# Spatial and temporal structure of choice representations in primate prefrontal cortex

**DOI:** 10.1101/595520

**Authors:** Sepp Kollmorgen, William T Newsome, Valerio Mante

## Abstract

Divergent accounts of how choices are represented by neural populations have led to conflicting explanations of the underlying mechanisms of decision-making, ranging from persistent, attractor-based dynamics to transient, sequence-based dynamics. To evaluate these mechanisms, we characterize the spatial and temporal structure of choice representations in large neural populations in prefrontal cortex. We find that the pronounced diversity of choice responses across neurons reflects only a few, mostly persistent population patterns recruited at progressively later times before and after a choice. Brief sequential activity occurs during a saccadic choice, but is entirely absent in a delay preceding it. The diversity of choice responses, which could result from almost-random connectivity in the underlying circuits, instead largely reflects the topographical arrangement of response-field properties across the cortical surface. This spatial organization appears to form a fixed scaffold upon which the context-dependent representations of task-specific variables often observed in prefrontal cortex can be learned.

## Introduction

Studies of decision-making in humans and animals provide a window onto the neural mechanisms underlying the interaction of sensory, cognitive, and motor processes, and have been singularly influential in shaping our understanding of neural computations across a variety of brain areas and species (Gold & Shadlen, 2007; Hanks & Summerfield, 2017; Schall, 2001; Shadlen & Kiani, 2013). A general framework explaining the neural mechanisms of decision-making, however, is yet to emerge, as studies on a variety of behavioral paradigms and animal models have led to rather different, and sometimes incompatible, explanations of how choices are generated and represented by neural circuits.

One line of research has focused on correlating the responses of single neurons in parietal and frontal areas to internal, decision-related variables like integration of evidence and confidence, which can be inferred from the animal’s choice-behavior (Hanks et al., 2015; Kepecs, Uchida, Zariwala, & Mainen, 2008; Kiani & Shadlen, 2009; Kim & Shadlen, 1999; Purcell, Schall, Logan, & Palmeri, 2012; Shadlen & Newsome, 2001; Yates, Park, Katz, Pillow, & Huk, 2017). Single-neuron responses can faithfully track these internal variables over durations lasting up to several seconds, suggesting that, at the level of the population, these variables are represented by patterns of activation that are low-dimensional and largely stable over time (Brody, Romo, & Kepecs, 2003; Ganguli et al., 2008; Machens, Romo, & Brody, 2005; Mante, Sussillo, Shenoy, & Newsome, 2013). Such stable dynamics can be explained by mechanistic neural models implementing attractor dynamics, i.e. patterns of population activity that persist in the absence of external inputs (Brody et al., 2003; Ganguli et al., 2008; Hopfield, 1982; Machens et al., 2005; Mante et al., 2013; Murray et al., 2017; Rolls, Loh, Deco, & Winterer, 2008; Seung, 1996; Wang, 2002).

A different line of research instead has emphasized the diversity and complexity of single-neuron responses observed in high-level association areas. Most neurons in these areas appear to represent not just one variable, but linear or non-linear mixtures of many behaviorally relevant variables (Hernandez et al., 2010; Jun et al., 2010; Machens, Romo, & Brody, 2010; Mante et al., 2013; Meister, Hennig, & Huk, 2013; Park, Meister, Huk, & Pillow, 2014; Parthasarathy et al., 2017; Raposo, Kaufman, & Churchland, 2014; Rigotti et al., 2013; Rishel, Huang, & Freedman, 2013; Singh & Eliasmith, 2006), resulting in a continuum of diverse single neuron responses that typically cannot be organized into distinct functional categories. Moreover, the representation of task variables by single neurons can be markedly transient, and become sustained only at the level of the entire population as a temporal sequence of activity patterns (Baeg et al., 2003; Crowe, Averbeck, & Chafee, 2010; Fujisawa, Amarasingham, Harrison, & Buzsaki, 2008; Goldman, 2009; Harvey, Coen, & Tank, 2012; Morcos & Harvey, 2016; Rajan, Harvey, & Tank, 2016; Scott et al., 2017). The representation of non-linear mixtures of variables, and the existence of long, apparently non-repeating temporal sequences, both suggest neural dynamics that are high-dimensional (Barak, Sussillo, Romo, Tsodyks, & Abbott, 2013; Goldman, 2009; Harvey et al., 2012; Rajan et al., 2016). Such high-dimensional dynamics are a critical feature of neural networks implementing reservoir computing (Jaeger & Haas, 2004; Maass, Natschlager, & Markram, 2002), which do not rely on attractor dynamics, but rather transform low-dimensional inputs (e.g. sensory evidence) into high-dimensional and task-dependent neural trajectories that can be easily read-out to produce the desired outputs (e.g. choice) (Buonomano & Maass, 2009; Jaeger & Haas, 2004; Maass et al., 2002; Rabinovich, Huerta, & Laurent, 2008).

Here we study the representation of choice in primate prefrontal cortex (PFC) during a decision-making task and in a delayed-saccade task, and determine whether the neural population responses in these tasks are more consistent with attractor dynamics, reservoir computing, or other computational schemes intermediate between these two extremes (Barak et al., 2013; Chaisangmongkon, Swaminathan, Freedman, & Wang, 2017; Rabinovich et al., 2008; Rajan et al., 2016). To distinguish between these possibilities, we focus on characterizing two properties of the population responses. First, we ask whether the representation of choice in the decision-making task is persistent or transient, both at the level of single neurons and of patterns of population activity. We find that single-neuron responses in both tasks are very diverse (Bruce & Goldberg, 1985; Chafee & Goldman-Rakic, 1998), and that profiles of activity across the population can be thought of as mixtures of only a few distinct population patterns, which have either mostly persistent or transient temporal dynamics (Machens et al., 2010; Mante et al., 2013; Singh & Eliasmith, 2006). Second, we determine to what extent choice responses in PFC are input and task-dependent, by asking if the responses in the decision-making task can be predicted based on responses in the delayed-saccade task. Critically, we find that the dynamics of single-neuron responses is largely preserved across tasks. Both findings are inconsistent with the high-dimensional, input-dependent dynamics expected from reservoir computing (Buonomano & Maass, 2009; Jaeger & Haas, 2004; Maass et al., 2002; Rabinovich et al., 2008).

Notably, much of the observed diversity of single-neuron dynamics in the decision-making task, which could result from almost-random connectivity in the underlying recurrent networks (Barak et al., 2013; Rajan et al., 2016; Rigotti et al., 2013; Sussillo, 2014), instead reflects the task-independent, topographical arrangement of single-unit response-field properties across the cortical surface. Such properties are not systematically probed by many current decision-making paradigms (Hanks & Summerfield, 2017; Shadlen & Kiani, 2013), which typically employ highly restricted subsets of possible spatial arrangements of sensory inputs and motor outputs, and thus by themselves seem inadequate to interpret the increasingly large population responses revealed by modern recording approaches (Cunningham & Yu, 2014).

## Results

### Behavioral task and neural recordings

We studied choice related neural activity in dorsolateral PFC of two macaque monkeys engaged in a two-alternative, forced-choice sensory discrimination task (Fig. 1; Supp. Fig. 1). The monkeys were trained to report the prevalent direction of motion in a random-dot stimulus (Britten, Shadlen, Newsome, & Movshon, 1992) (e.g. left vs. right) with a saccade to one of two choice targets (Fig. 1a), and were rewarded for correct choices. While the monkeys performed this task, we recorded single- and multi-unit activity with a multi-electrode array (96 channels) chronically implanted in the left pre-arcuate gyrus (Kiani, Cueva, Reppas, & Newsome, 2014; Kiani et al., 2015; Schall, 1997) (area 8Ar, Supp. Fig. 2). In each experimental session, we recorded the simultaneous activity from 185±43 units per session (mean±std) in monkey T and 241±46 in monkey V, for a total of 185 and 184 sessions distributed over 67 and 62 days, with a median session duration of 293 and 173 trials. Below we focus on data from monkey T, whereas the largely analogous data from monkey V is shown in the supplementary material.

**Figure 1.**
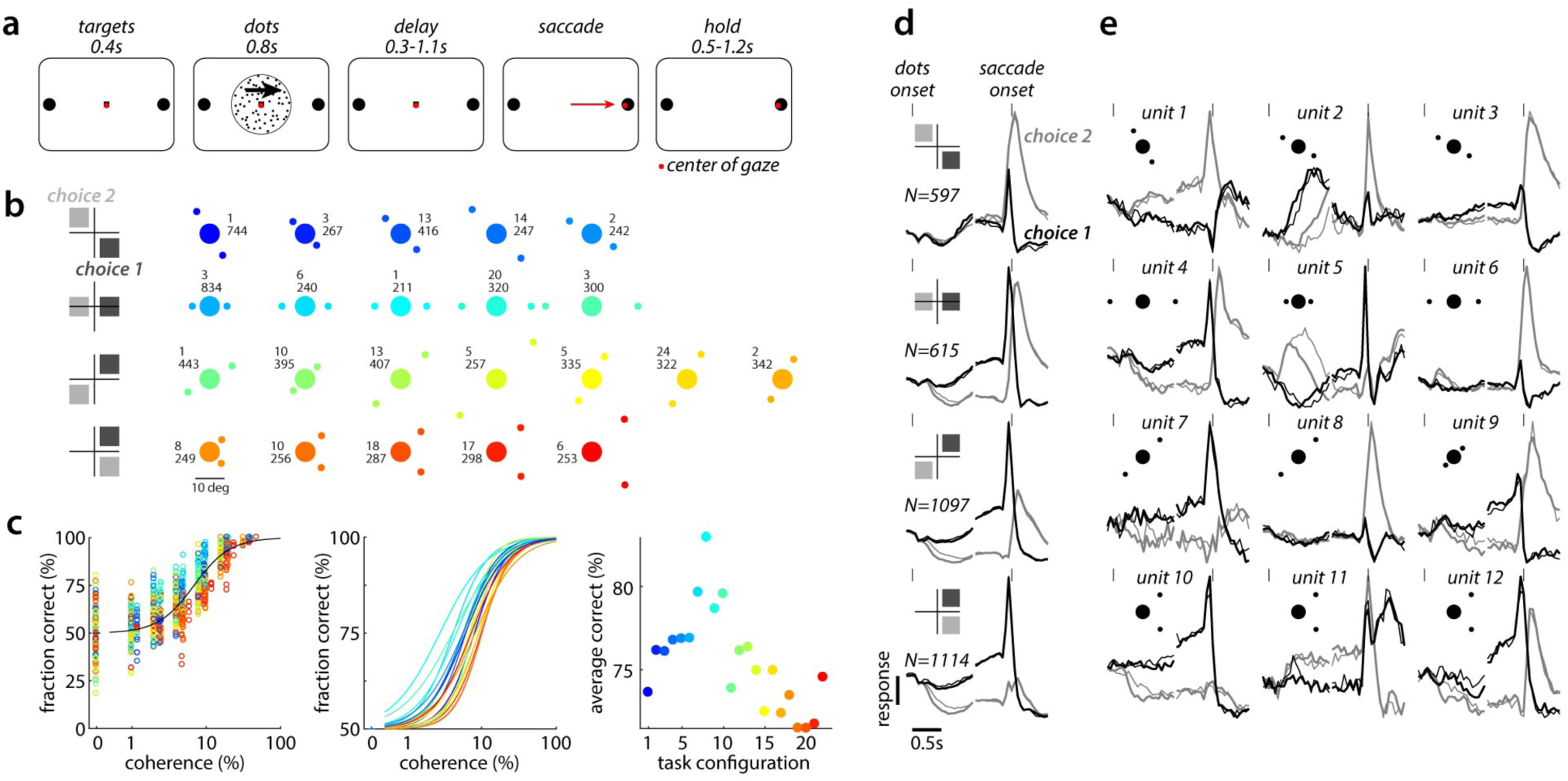
Behavioral task and neural responses in PFC. **a**, Behavioral task. Monkeys kept their center of gaze (red dot) on a fixation point, indicated the perceived direction of motion of a random-dots stimulus with a saccade to one of two targets, and received a reward (on correct trials) after briefly fixating on the chosen target (hold period). **b**-**e**, Behavioral performance and neural responses for monkey T. **b**, Target configurations. The random dots were always centered on the fixation point, while the two targets were arranged in different configurations across sessions (insets: number of sessions, top; average number of behavioral trials per session, bottom; target diameter is not shown to scale). We grouped the 22 different target-configurations into 4 “task-configurations” (rows). **c**, Behavioral performance, same colors as in **b**. Left panel: fraction correct as a function of motion strength (coherence) and configuration. Middle: Fits of a behavioral model for each configuration, based on the data in the left panel. Right: average performance for each configuration, as estimated from the fits (middle) over a set of coherences common to all configurations. **d**, Average responses over units recorded in one of the four task-configurations in **b**. Normalized, de-noised responses are aligned to dots onset and saccade onset (tick marks, top) and averaged based on the location of the chosen target (choice 1, black; choice 2, gray, defined as in **b**) and outcome (correct, thick; error, thin). Responses were de-noised with Targeted Dimensionality Reduction (Supp. Fig. 3). Here only the 10% most choice-predictive units in each task configuration are averaged. **e**, Example de-noised responses from individual units, selected to illustrate the range of unit responses in the population. Units were recorded in different target configurations (insets) and at different cortical locations (not shown). These units are shown also in Fig. 6b (red crosses).

**Figure 2.**
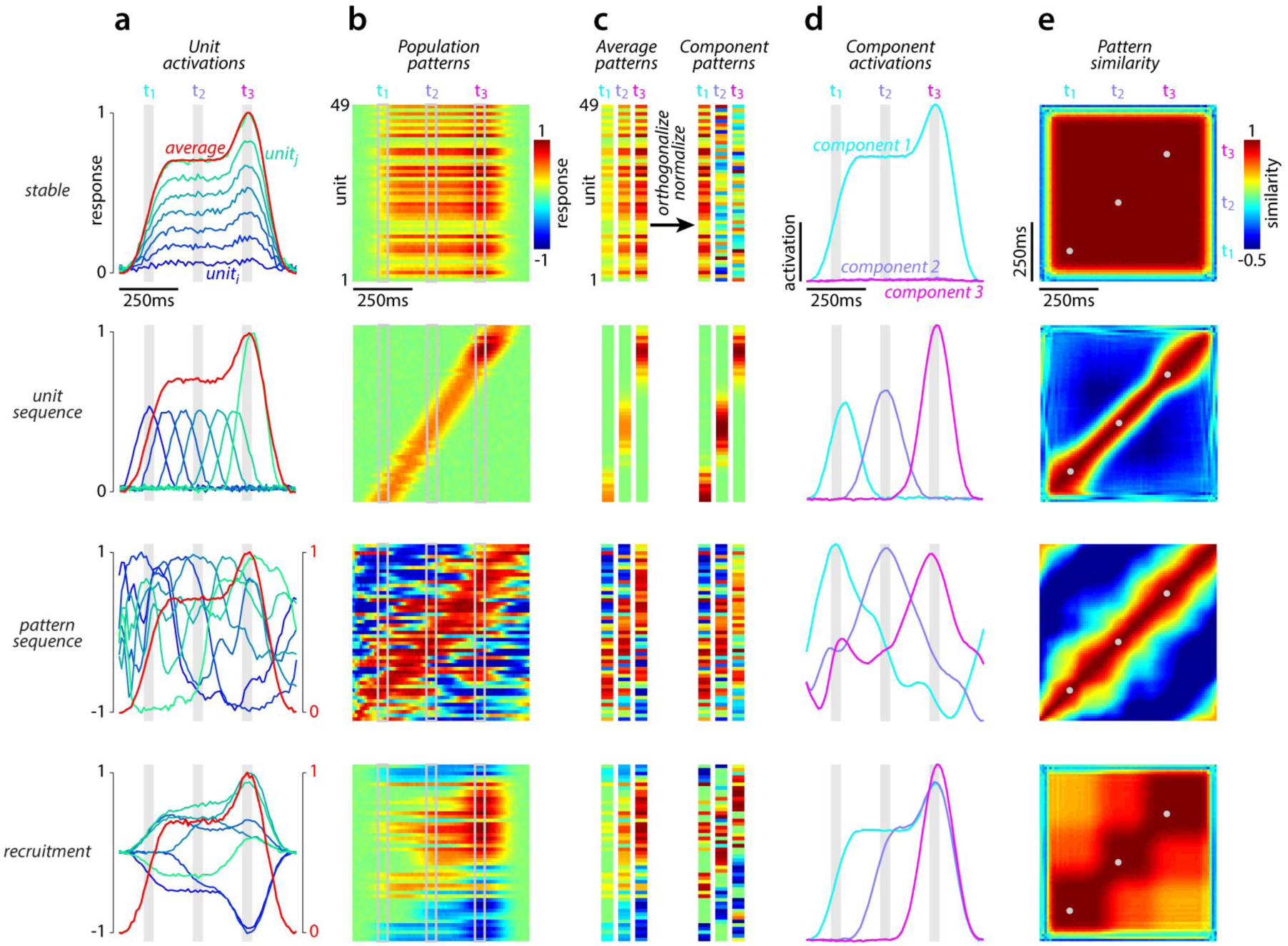
Different hypothesized representations of choice in the population. Each row illustrates the temporal dynamics of single unit and population activity for four idealized representations of choice: (1) a stable representation (first row from top); (2) a sequence across units (second row); (3) a sequence across patterns (third row); (4) the serial recruitment of distinct choice patterns (fourth row). Representations 1, 2, and 4 were constructed by hand. Representation 3 are simulations of a recurrent neural network, trained to produce the population average response of representation 2. **a**, Responses of representative single units (blue to green; analogous to choice 1 responses in Fig. 1e) and normalized population average (red; analogous to choice 1 response in Fig. 1d; responses to choice 2 are set to zero). In the third and fourth row, single-unit and population average responses are shown with respect to different baselines (compare left and right vertical axes). **b**, Responses of many single units, ordered along the vertical axis based on the time of their peak-response. Activity along any vertical line in each plot describes the population activity pattern at the corresponding time. For the stable and recruitment representations, the exact arrangement of units along the vertical axis is strongly affected by noise. For example, in the case of recruitment the blue bands at the top and bottom of the plot correspond to units whose largest firing rate is close to zero, occurring either at the beginning or end of the trial depending on the noise. **c**, Definition of component patterns. We first averaged population patterns over time within three non-overlapping time windows (t_1_, t_2_, and t_3_; gray squares in **a**, **b**, and **d**). The resulting average patterns (left) are orthogonalized and normalized to obtain the component patterns (right). **d**, Definition of component activations. Each component activation is obtained by taking the “running” dot-product of the corresponding component pattern with the population patterns in **b**. The component activations measure how much each component pattern contributes to the population pattern at any given time. **e**, Similarity of population patterns across time. For every pair of times, similarity is computed as the dot-product of the (normalized) population patterns in **b** at the corresponding times. The center of the time windows used to define the component patterns are indicated by gray dots.

Each behavioral trial consisted of a stereotyped sequence of events (Fig. 1a). The monkeys initiated each trial by looking at a fixation point. After a short delay, the two choice targets appeared, followed by the random-dot stimulus of fixed duration (0.8s). The offset of the random-dots initiated a delay-period of random duration (0.3-1.1s). The delay-period ended with the disappearance of the fixation point, which instructed the monkey to quickly indicate its choice with a saccade to one of the targets. The saccade was followed by a hold period of random duration (0.5-1.3s), during which the monkey had to maintain fixation on the chosen target until a feedback tone was played (correct or wrong) and the reward delivered.

The location of the two choice targets was varied across sessions to obtain extensive coverage of both visual hemi-fields, while the random-dots stimulus was always shown at the center of gaze (Fig. 1b). The monkeys were highly proficient at the task in all target configurations (Fig. 1c), though performance was noticeably better for the configurations that were used most often during training (horizontal target arrangement).

The activity of a large fraction of recorded units was related to the monkey’s upcoming choice at some point during the trial (64±10% and 43±10% in monkeys T and V, p=0.05 corrected for multiple comparisons; null distribution based on random permutations of all trials, see Methods). For each unit, here we separated trials based on the combination of choice (choice 1 vs. choice 2) and outcome (correct vs. error), resulting in four condition-averages for each unit (Fig. 1d,e; Supp. Fig. 3). In most task configurations, we define saccades to the target in the right visual hemi-field (contralateral to the recording array) as choice 1, and saccades to the other target as choice 2. When both targets are in the right hemi-field, choice 1 is defined as the saccade to the upper hemi-field (Fig. 1b). With this definition, population-average responses during the delay period tend to be larger for choice 1 than choice 2 (Fig. 1d). Below we use such condition averages to characterize the responses of individual units and of the population. Trial-by-trial variability of responses within a condition, which may further constrain the nature of the underlying neural processes (Bollimunta, Totten, & Ditterich, 2012; A. K. Churchland et al., 2011; M. M. Churchland et al., 2010; Latimer, Yates, Meister, Huk, & Pillow, 2015; Morcos & Harvey, 2016; Seidemann, Meilijson, Abeles, Bergman, & Vaadia, 1996) is mostly not considered here.

**Figure 3.**
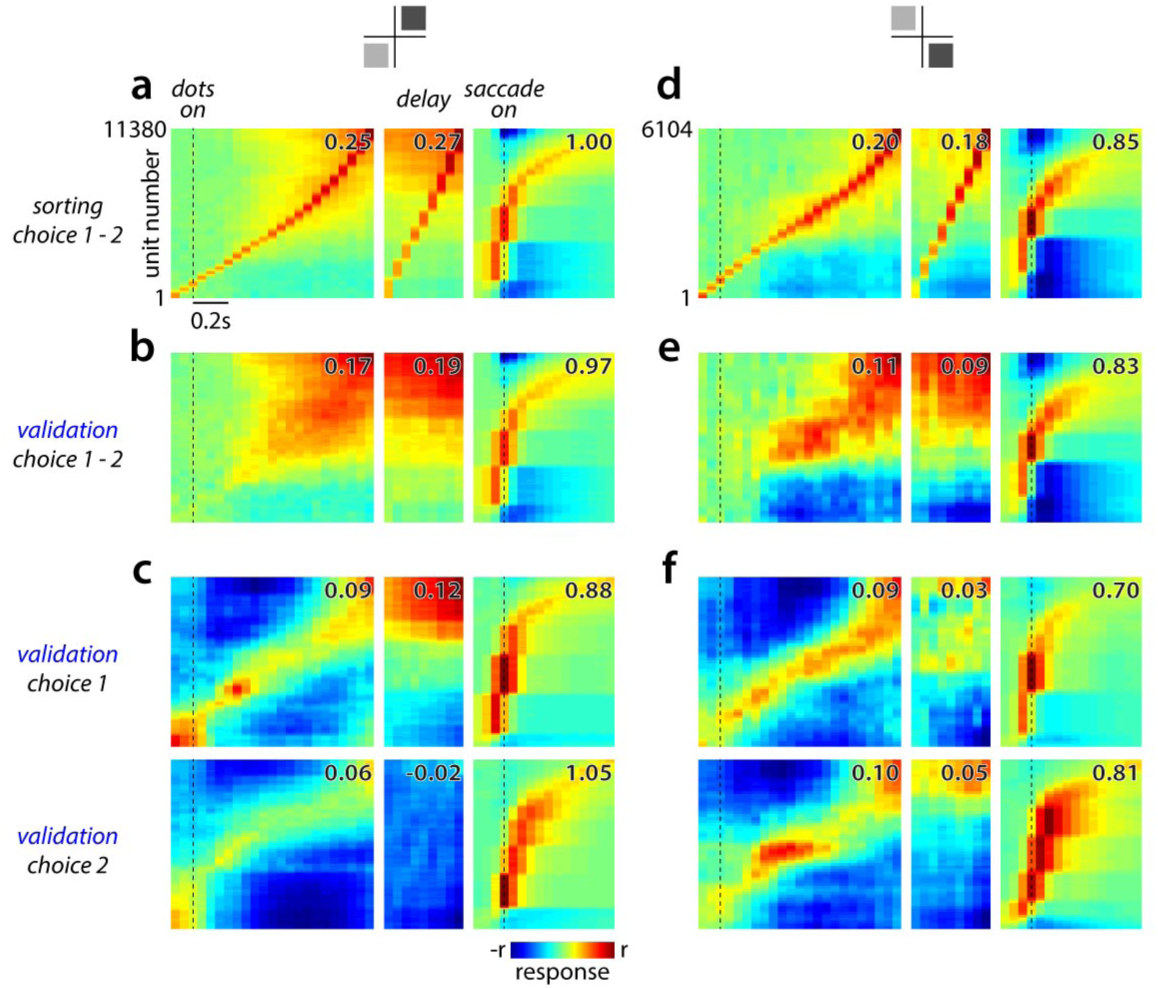
Dynamics of unit responses in PFC. Average de-noised responses for all units recorded in two task-configurations (**a**-**c** and **d**-**f**, top insets as in Fig. 1b), sorted by peak-time (monkey T), analogous to Fig. 2b. Responses are aligned to dots onset (left sub-panels) or saccade onset (middle and right, showing contiguous times in the delay and saccade epochs). Peak-time is determined separately for the dots, delay, and saccade epochs. Because of the large number of units in each configuration, after arranging the sorted responses in a two-dimensional image as in **a**, we smooth (moving square filter, width 250 rows) and subsample (every 100th row) the responses along the vertical axis. The numbers in the top-right of each panel indicate the peak response in each resulting smoothed image (colorbar, r). **a**, Choice-related activity, defined as the difference between condition-averaged responses for choice 1 and choice 2 trials (correct only). Averages are computed based on a randomly selected half of the trials (sorting set). Units are ordered along the vertical axis based on the time of maximal choice-related activity (early, bottom; late, top). **b**, Choice-related activity based on the trials not used in **a** (validation set). The order of units along the vertical axis is maintained from **a**. **c**, Average responses to choice 1 (top) and choice 2 (bottom), sorted separately based on the respective peak-times. Only responses from the corresponding validation sets are shown. **d**-**f**, Analogous to **a**-**c**, for a different task configuration.

The recordings reveal a considerable diversity of responses across individual units. Choice-related responses do not occur simultaneously across all units, but rather can occur at different times in different units (Fig. 1e). Indeed, the average response over a large number of units (Fig. 1d) resembles the activity of some individual units but not that of others (Fig. 1e). The extent of heterogeneity across the population, however, is difficult to assess based on such anecdotal examples, which may represent outliers of a rather homogeneous population. To obtain a more complete picture of the diversity of choice responses, below we characterize the responses of all units in the population.

### Hypothesized choice representations

We start by considering four simulated neural populations that represent choice in fundamentally different ways (Fig. 2). We will use these simulated responses as four hypotheses that can be compared to the responses measured in PFC, and to illustrate several complementary analysis approaches that together can distinguish between these hypotheses. The simulations are not meant to faithfully reproduce the diversity of responses observed in the recorded units, but rather represent idealized implementations of different possible choice representations.

The four simulated populations are shown in the four rows in Fig. 2. In the first population, choice is represented by a single, stable pattern of activation, meaning that the response of all single neurons in the population are essentially scaled versions of each other (Fig. 2a, *stable*). In the second population, each neuron responds only transiently, at a consistent time during the trial, with different neurons representing choice at different times during the trial (Fig. 2a, *unit sequence*). In the third population, the representation of choice is passed sequentially along a chain of activation patterns, with single neurons showing diverse, multi-peaked responses (Fig. 2a, *pattern sequence*). Finally, the fourth population contains patterns of activation that are persistent, but are recruited at progressively later times during the trial, again resulting in diverse single neuron responses (Fig. 2a, *recruitment*). Experimental evidence for most of these representations has been previously reported (e.g. stable (Ganguli et al., 2008; Machens et al., 2005; Mazurek, Roitman, Ditterich, & Shadlen, 2003; Wang, 2002), sequence of units (Baeg et al., 2003; Fujisawa et al., 2008; Harvey et al., 2012; Morcos & Harvey, 2016; Rajan et al., 2016), sequence of patterns (Bollimunta et al., 2012; Goldman, 2009; Wehr & Laurent, 1996), and combinations thereof (M. M. Churchland et al., 2012; Kaufman, Churchland, Ryu, & Shenoy, 2014)).

A first approach to distinguishing some of these representations relies on visualizing the activity of all units after sorting them by the time of peak-activation (Fujisawa et al., 2008; Harvey et al., 2012; Morcos & Harvey, 2016) (Fig. 2b). When the population response is organized as a sequence of units, these plots reveal a prominent diagonal band (Fig. 2b, *unit sequence*), which corresponds to the “wave” of activity traveling across the population. This diagonal band is absent for a stable representation (Fig. 2b, *stable*) and for recruitment (Fig. 2b, *recruitment*). A similar band can be observed for a sequence of patterns— the band however is less prominent, as individual units that are active more than once over the course of the trial contribute to “off-diagonal” responses (Fig. 2b, *pattern sequence*).

A second approach relies on characterizing the temporal dynamics of population patterns, rather than single neurons (Fig. 2c,d). The population activity pattern at a given time during the trial is given by the responses along the corresponding vertical line in Fig. 2b. Differences between the representations are revealed by asking how long the pattern observed at any given time persists in the population response. To answer this question, we first define three *average patterns* by averaging responses over time within three distinct temporal windows (Fig. 2c, left; t_1_-t_3_ windows: gray rectangles in Fig. 2a, b, and d). We then obtain three *component patterns* by orthogonalizing the three average patterns (Fig. 2c, right). The orthogonalization ensures that each component pattern describes features of the population response that are not already captured by the previous component patterns. Finally, we compute *component activations* (Fig. 2d) by taking the dot product of each component pattern (Fig. 2c, right) with the population patterns at each point in time (Fig. 2b). These steps amount to a dimensionality reduction approach (Briggman, Abarbanel, & Kristan, 2005; M. M. Churchland et al., 2012; Cunningham & Yu, 2014; Kobak et al., 2016; Laurent, 2002; Mante et al., 2013; Yu et al., 2009), as each component pattern corresponds to a dimension in state-space, and the activations capture the contribution of the corresponding dimension to the population response.

The component activations reveal many defining features of the underlying representations. For the stable representation, the first component pattern (Fig. 2c, stable; right, t_1_) is persistently active (Fig. 2d, stable; component 1) beyond the time window used to define that component (Fig. 2d, stable; gray bin at t_1_). The second and third component patterns (Fig. 2c, stable; right, t_2_ and t_3_) essentially reflect noise, and show very small activations (Fig. 2d, stable; components 2 and 3). For a sequence of units or patterns, the component activations are transient (Fig. 2d, unit sequence and pattern sequence), and large only during the time window used to define the corresponding component pattern, indicating that the population continuously undergoes smooth transitions from one activity pattern to another. In the case of recruitment, several components can show persistent activation, each starting from the time used to define the corresponding pattern (Fig. 2d, recruitment).

A third approach to comparing representations relies on computing the similarity between activity patterns measured at different times in the trial (i.e. activity along any possible vertical line in Fig. 2b). This approach does not provide all the insights revealed by the component activations, but has the advantage of not depending on a particular choice of average patterns as in Fig. 2c. For any pair of times in Fig 2b, we define similarity as the correlation between the corresponding activity patterns. We plot the full set of similarities as the matrices in Fig. 2e, with each point in the matrix corresponding to a pair of time bins in Fig 2b. Since patterns extracted at nearby times are similar, similarity is largest close to the positive diagonal (Fig. 2e; from bottom-left to top-right). The different persistence of responses in the stable, sequential, and serial recruitment representations is reflected by how far these large similarity values extend away from the diagonal, i.e. to more dissimilar times (Fig. 2e).

### Dynamics of single unit responses

Below we use these three approaches to characterize the dots-task responses in PFC. As for the simulated responses (Fig. 2b), we first sort units by the time of peak-activation (Fig. 3; Supp. Fig. 4). Since for some units the response for a given choice, or the difference in the response to the two choices, shows peaks at multiple times (e.g. Fig. 1e, units 2, 5, or 6) we sort the units repeatedly based on the responses in three distinct task epochs, namely the dots-period (Fig. 3a,d; left panel), the delay period (middle), and the saccade/hold periods (right). To exclude contributions from trial-by-trial variability to these plots, we applied a cross-validation procedure. For each unit, we randomly assigned trials into a *sorting* or a *validation* set—we used responses from the sorting set to order units based on their peak activation-time (Fig. 3a,d) and responses from the validation set to evaluate the existence of a sequence (Fig. 3b,e). The ordering of units along the vertical axis across the resulting two plots is preserved, but is entirely determined by responses in the sorting set.

**Figure 4.**
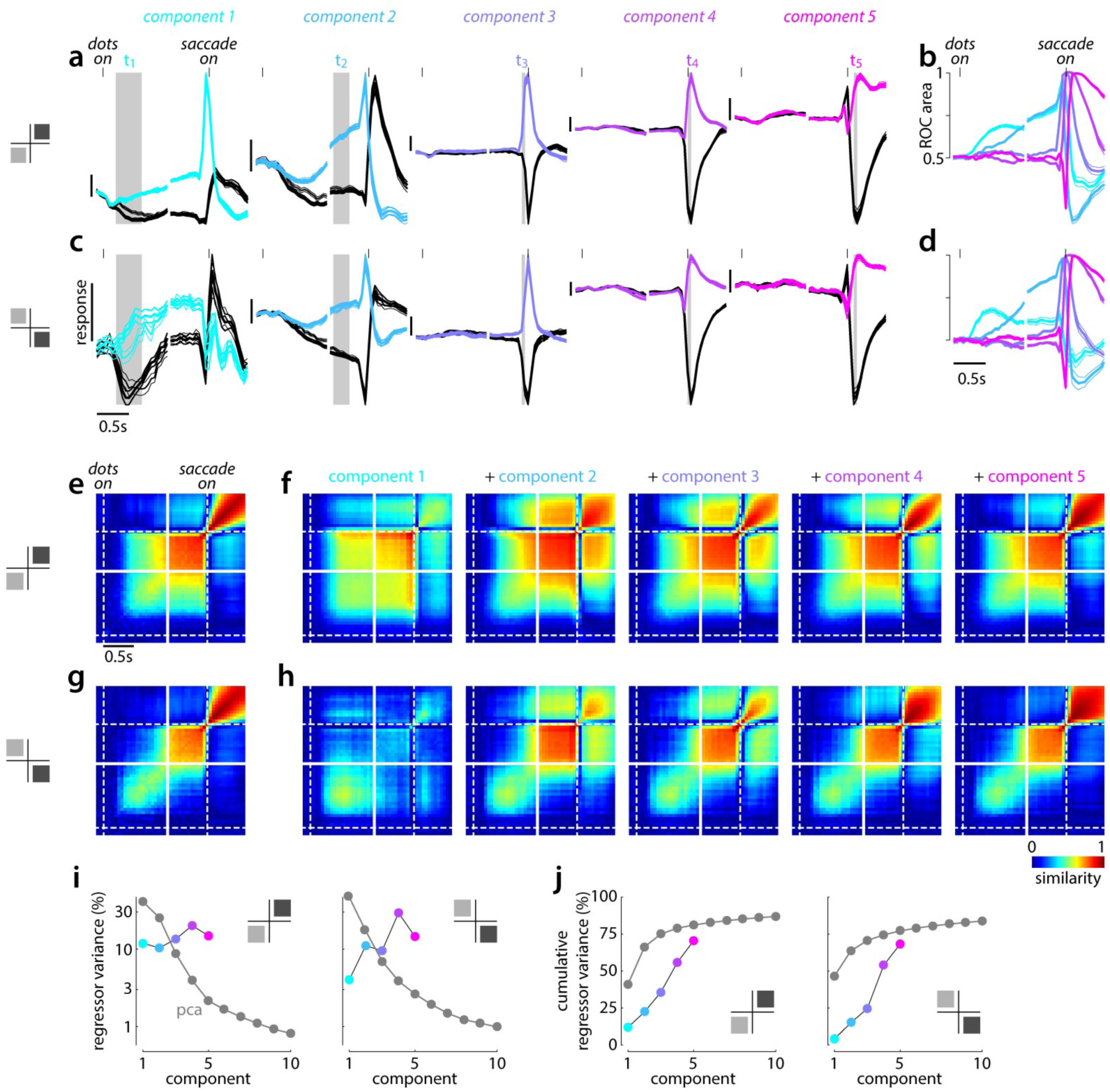
Dynamics of population activity patterns in PFC. Data from monkey T. **a**, Component activations for choice 1 (colored) and choice 2 (black) on correct (thick) and incorrect (thin) trials (analogous to Fig. 2d). The five components were estimated separately for each recording session from population patterns defined at the times indicated by the corresponding vertical gray stripes, and the resulting component activations were averaged over all sessions belonging to a given task-configuration (inset on the left). Error bars are the standard error of the mean. **b**, Separation between choice 1 and choice 2 component activations, measured as area under the ROC curve for the corresponding distributions of trial-by-trial activations. **c**,**d**, Same as **a**,**b**, for a different task configuration (left inset, as in Fig. 1b). **e**-**h**, Measured and predicted similarity of choice predictive patterns estimated at different times in the trial (analogous to Fig. 2e). Similarity is computed separately for each session, and averaged across sessions from the same task configuration (inset, left). Similarity is cross-validated (see methods), and thus can be smaller than 1 even for patterns extracted at the same time (points on the diagonal from bottom-left to top-right). **e**, Measured similarity between choice-predictive population patterns for one task configuration. **f**, Predicted similarity, based on population responses reconstructed with increasing numbers of choice-related component patterns (from component 1 only, to all five components; left to right). Substantial random noise was added to each unit’s response in the reconstruction. Unlike in Fig. 2e top, similarity thus varies across pairs of times even when only component 1 is used (left-most panel) and reflects the difference in magnitude between choice 1 and choice 2 responses for component 1 (left-most panels in **a**,**c**). **g**,**h**, Same as **e**,**f**, for a different task configuration (left inset; as in Fig. 1b). **i**, Variance in the choice-related population patterns explained by the 5 component patterns (colored) or by 10 principal components. Task configurations as in **a** (left) and **b** (right). **j**, Cumulative variance explained, computed from the plots in **i**. In **i** and **j**, standard error of the mean over sessions is smaller than the symbols.

The resulting cross-validated plots show little evidence of sequences throughout much of the trial (Fig. 3b,e). Here, we generated these plots directly for the choice–related activity, measured as the difference between choice 1 and choice 2 responses. A clear, but brief sequence of units can be observed starting from about 100ms before saccade onset to about 400ms after (Fig. 3b,e, right). During the dots period, responses also peak at different times in different units (Fig. 3b, left), although peak-times might not be distributed evenly across the population in all task configurations, but rather may fall into two largely separate time windows, early and late after dots onset (Fig. 3e, left). During the delay period, we find no evidence of a sequence irrespective of task configuration (Fig. 3b,e; middle panels). Interestingly, a more prominent sequence can be observed during the dots period if choice 1 and choice 2 responses are considered separately (Fig. 3c,f). These sequences appear to reflect activity that is common to both choices, and is thus less prominent when considering the difference between choice 1 and choice 2 activity (Fig. 3b,e).

### Dynamics of population activity patterns

To characterize the dynamics of population activity patterns (Fig. 4; Supp. Fig. 5), we restrict the analysis to choice-related activity in the population, by considering only activity patterns that are modulated by choice. We use linear regression to extract the pattern of population activity that best predicts the upcoming choice at any given time during the trial (Mante et al., 2013), define component patterns (as in Fig. 2c) by averaging the extracted choice predictive patterns within different time-windows in the trial (gray stripes in Fig. 4a,c, t_1_ to t_5_), and then compute the corresponding component activations as the dot product between the component patterns and the population activity (Fig. 4a,c). As for single unit responses (Fig. 1d), we computed condition-averaged component activations based on choice and trial outcome (Fig. 4a,c).

**Figure 5.**
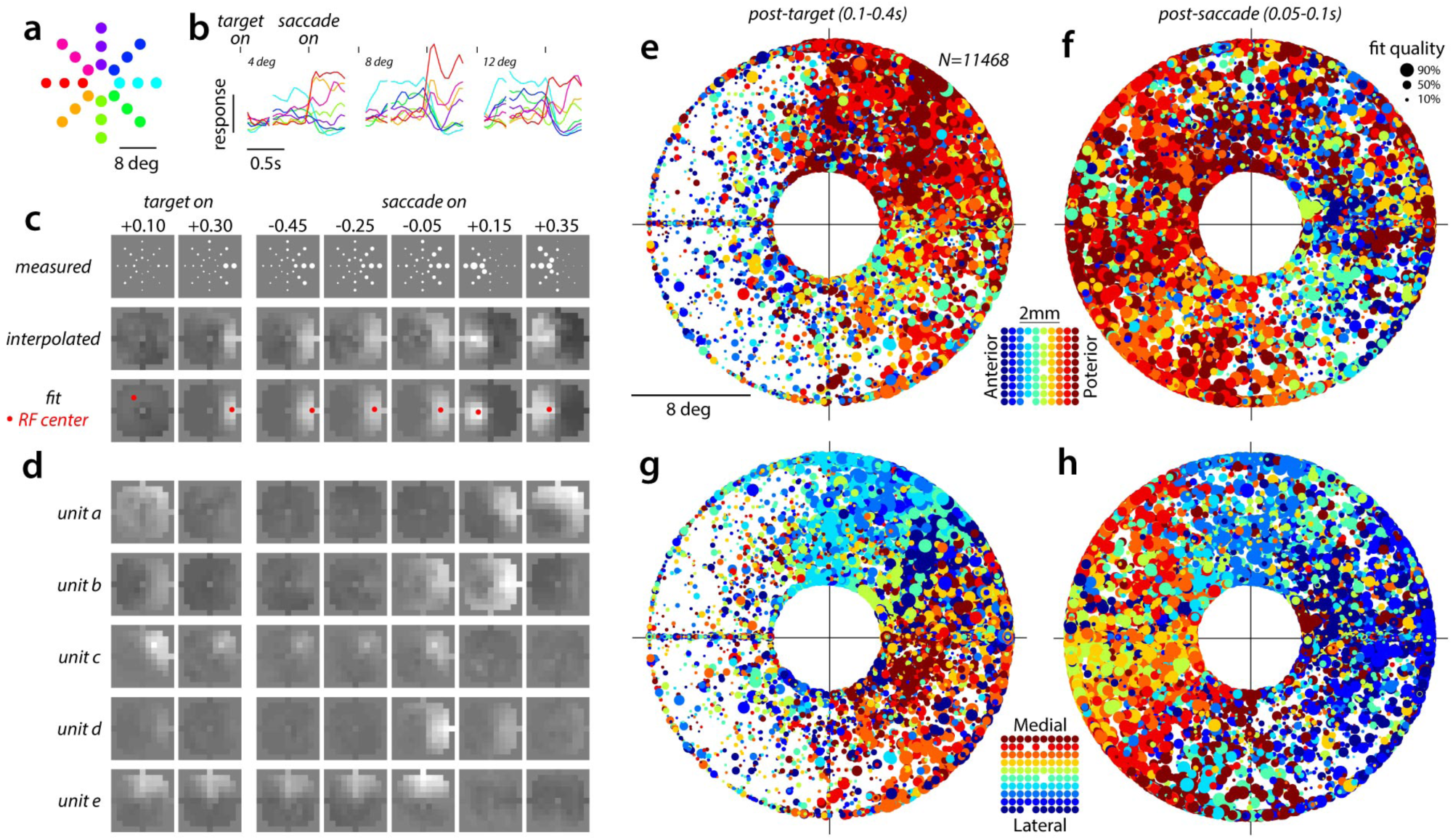
Response-field properties in a delayed-saccade task. Data from monkey T. **a**, Locations in visual space of all saccade targets used in an example session of the visually guided, delayed-saccade task. **b**, Normalized average responses during the delayed-saccade task for an example unit. Same target locations as in **a**, varying in radius (left to right panels) and angular location (color). Responses are aligned to target onset and saccade onset (tick marks, top). **c**, Responses of the unit in **b**, replotted at discrete times during the trial (relative to target and saccade onset, top numbers in seconds) at the location of the corresponding target (white circles, radius proportional to response). We estimated the time-varying response-field of a unit by either interpolating the responses at regular grid locations extending over the possible target locations (middle) or by fitting a response-field model to the responses (bottom). From the model fits we extracted the response-field center (i.e. the peak location, red points) at each time. **d**, Interpolated response-fields for five example units, showing a diversity of visual, delay-period, movement, and hold-period responses. **e**, Response-field centers for all units recorded in the delayed-saccade task (circles) estimated from responses obtained 0.1-0.4s after target onset. Each circle marks the location of a response-field center in the visual field. Circle radius is proportional to the quality of the underlying fit (percentage of variance explained, legend) and can be interpreted as the strength of spatial tuning (strong vs. weak; large vs. small circles). Units are colored based on their anterior-posterior location along the electrode array (inset) and are plotted in random order. **f**, same as **e**, for responses at the beginning of the hold period (0.05-0.1s after saccade onset). **g**-**h**, same data as in **e**-**f**, but with units colored based on their medio-lateral location on the array (inset).

This approach reveals both persistent and transient choice components (Fig. 4a,c), which are predictive of choice at different times during the trial (as quantified by the area under the ROC curve, Fig. 4b,d). A first choice-related pattern (component 1) emerges shortly after dots-onset, is largely persistent across the dots and delay periods, peaks around the time of saccade initiation, and shows inverted selectivity for choice after the saccade. A second persistent pattern (component 2) emerges later during the dots presentation, becomes increasingly choice-predictive during the delay period, and also shows inverted selectivity after the saccade. A third pattern (component 3), defined at the time of saccadic initiation, is instead mostly transient. Two additional patterns defined after saccade initiation (components 4 & 5) peak at increasingly later times during the hold period prior to reward (Fig. 1a), as expected by sequential activation, but are also largely persistent throughout the hold period.

The component activations overall mirror the characterization of the dynamics at the level of individual units (Figs. 1e and 3) and suggest a representation with prominent serial recruitment of components before the saccade (e.g. compare components 1-3 in Fig. 4a,c to Fig. 2d, *recruitment*), fast sequential encoding around the time of the saccade (e.g., choice 1 responses peak at increasingly later times for components 1-5 in Fig. 4a, as for components 1-3 in Fig. 2d, second and third rows), and a slow sequence and possibly recruitment after the saccade (e.g. choice 1 responses for components 4,5 in Fig. 4c & Supp. Fig. 5a,c). This conclusion is further supported by the similarity matrices for PFC responses (Fig. 4e,g— same format as Fig. 2e), which show periods of persistent, or slow, dynamics during the delay and hold periods (i.e. large similarities away far from the diagonal), and transient dynamics around the time of the saccade (large similarities only along the diagonal). The time-course of similarity, however, is not a fixed property of the neural population, but rather depends on task-configuration—in some configurations (Fig. 4e) the early choice signals appear to be more persistent than in others (Fig. 4g), again in qualitative agreement with the other analyses (compare Figs. 3b and e; and Figs. 4a and c, component 1).

Notably, these five component activations provide a largely complete description of choice responses in single sessions of the dots-task. To illustrate this point, we reconstructed the response of every recorded unit as a weighted sum of the component activations (Kobak et al., 2016), with weights varying across components and units (the weights are discussed in more detail below, see Fig. 5g). The observed diversity of single-unit responses (e.g. Fig. 3) reflects differences in the relative weights of the five components across units. We then computed similarities on the reconstructed population responses (Fig. 4f,h) as we did for the recorded responses (Fig. 4e,g). When each unit’s response is reconstructed based on only the first component, the match between predicted (Fig. 4f,h; left-most panel) and measured (Fig. 4e,g) similarities is poor. However, as more and more components are added (Fig. 4f,h; additional panels) the match progressively improves, and becomes very good when all five components are used to reconstruct the responses (Fig. 4f,h; right-most panel; correlation between measured similarity in Fig. 4e,g and predicted similarity in the right-most panels of Fig. 4f,h is R=0.993 and R=0.991, for the two task configurations). Additional component patterns that can be defined from the responses account only for a small fraction of the variance in the choice-predictive activity (Fig. 4i,j; Supp. Fig. 4i,j).

### Measuring response-fields with a delayed-saccade task

Having characterized the temporal dynamics of responses in a decision-making task (Figs. 3,4) we aim to relate these responses to those in a visually-guided, delayed-saccade task (Fig. 5; Supp. Fig. 6). If PFC were implementing strongly input-dependent dynamics akin to reservoir computing, single unit responses in the two tasks can be expected to be substantially different (Buonomano & Maass, 2009; Jaeger & Haas, 2004; Maass et al., 2002; Morcos & Harvey, 2016; Rabinovich et al., 2008).

**Figure 6.**
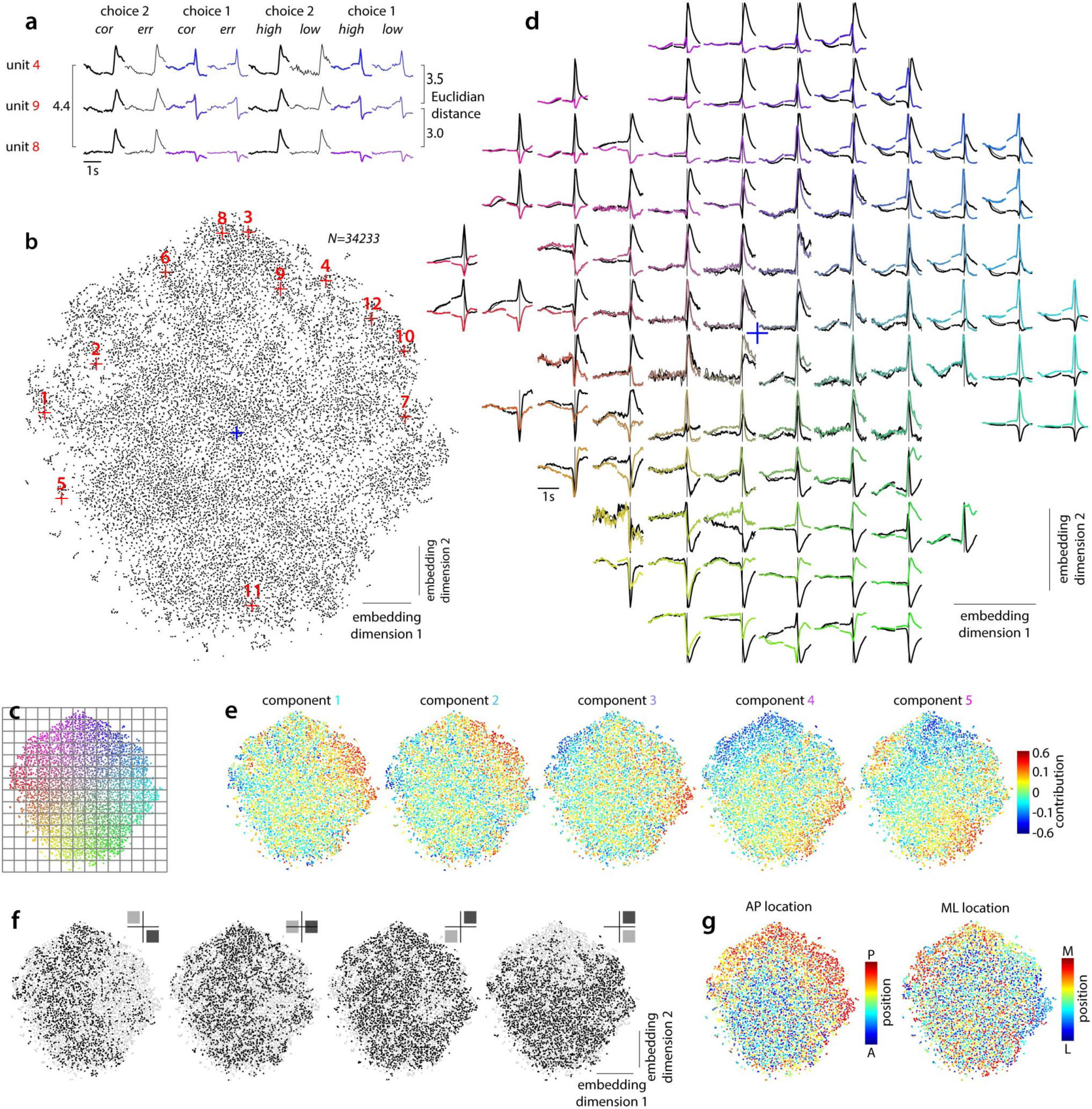
Global structure of single-unit responses in the dots-task. Data from monkey T. **a**, Three example units from Fig. 1e. Each individual unit is characterized by the averaged, de-noised responses on eight conditions defined based on choice (1 or 2), outcome (correct, cor; error; err), and motion coherence (high and low, only for correct trials). Here responses are aggressively de-noised by reconstructing each unit’s activity based on the five choice components estimated in the corresponding recording session (averages across sessions shown in Fig. 4a,c). For each unit, we concatenate all condition-averages into a single vector, and then compute Euclidian distances between vectors. **b**, Two-dimensional, non-linear embedding of all units in the population obtained with t-SNE. Each point corresponds to a unit. The arrangement of units within the two-dimensional embedding space optimally maintains nearest-neighbor relations between units in the high-dimensional space, defined based on distances computed as in **a**. The red crosses indicate the twelve example units in Fig. 1e. **c**,**d**, Average unit responses for nearby units at different locations along the embedding dimensions. We overlaid a regular 12×12 grid over the embedding space (panel **c**) and averaged the responses of all units falling within a given grid-square to obtain the responses at the corresponding locations in **d**. Only the first four conditions in panel **a** are shown. Choice 1 averages are colored based on the radial and angular position of the corresponding units in the embedding space (panel **c**). The blue cross marks the origin of the embedding space, as in **b**. **e**-**g**, Neural, task, and cortical factors contributing to the diversity of single-unit responses. The arrangement of units in each panel is identical to **b**. In **e** and **g** units are plotted sequentially in a random order. **e**, The contributions of the five component activations to the responses of each unit. Reconstructed unit responses as in **a** are obtained as a weighted sum of the component activations (Fig. 4a,c) from the corresponding recording session, with the weights given by the component contributions shown here. The relative weights of the five components differ across units, resulting in the diversity of unit responses in **d**. **f**, Units recorded in a given task configurations (black points; task-configurations as Fig. 1b) overlaid over all units in the population (gray points). Some unit responses (i.e. embedding locations) are not observed in all task configurations. **g**, Cortical location of all the units in the population. AP, anterior-posterior axis along the electrode array; ML: medio-lateral axis.

In the delayed-saccade task, the monkeys were rewarded for making a saccade to a single target, whose location was varied across trials to obtain extensive coverage of the visual field (Fig. 5a). Only one target was presented on each trial, and no random-dots were shown, but the timing of task events was otherwise analogous to the dots-task (Fig. 1), in particular with respect to the durations of the delay and hold periods.

For any given unit, the condition-averaged responses for each target location (Fig. 5b) are best visualized as a time-dependent response-field, representing the spatial tuning of visual, delay, movement, and hold period activity (Fig. 5c). At each time in the trial, we obtain smooth response-fields from the condition-averaged responses (Fig. 5c, top) either through linear interpolation (Fig. 5c, middle) or by fits of a simple descriptive model that assumes separable tuning for eccentricity and angular location (Bruce & Goldberg, 1985) (Fig. 5c, bottom). We use the fits to estimate the response-field center at each time, i.e. the spatial location of the peak response at that time (Fig. 5c, red dots; Fig. 5e-h).

The response-fields can be widely different across different units, both in terms of spatial organization and temporal dynamics (Fig. 5c,d). A complete characterization of all observed response-fields is beyond the scope of this study, but one can easily show that the response-field properties are not randomly distributed across cortical locations. To visualize the underlying topography, we relate the cortical location of each unit to the visual location of its response-field center after target onset (Fig. 5e,g) and after saccade onset (Fig. 5f,h). We show each response-field center twice, either colored based on the anterior-posterior (Fig. 5e,f) or the medio-lateral location (Fig. 5g,h) of the array electrode where it was recorded. The majority of units responding to the target onset are most active when the target appears in the right (contralateral), upper visual quadrant (Fig. 5e,g). These units are mostly recorded in posterior locations on the array (Fig. 5e, red colors), and their angular preference varies based on medio-lateral location—units from lateral array locations prefer targets in the upper-right quadrant and units from medial locations prefer the lower-right quadrant (Fig. 5g, blue vs. red colors). Post-saccadic responses are more broadly distributed across the array, but again the target preferences map regularly onto the cortical surface (Fig. 5f,h). Post-saccadic responses to targets in the left (ipsilateral) and right (contralateral) hemifield mostly occur at posterior and anterior locations, respectively (Fig. 5f, red vs. blue colors) while angular preference depends on medio-lateral location (Fig. 5h).

Relating these response-fields to the unit responses in the dots-task (Figs. 3,4) is challenging, because we could not without a doubt identify units that were recorded in both tasks (see Methods, Neural recordings). A direct comparison of responses in the two tasks is further complicated because on any given trial of the delayed-saccade task, only one response target, and no random-dots, appear on the screen. The responses in the two tasks thus can be expected to be different even if fixed, task-independent response-fields were to explain the responses in both settings.

To overcome this challenge, instead of comparing responses in the two tasks unit-by-unit, we compare them at the level of the population, focusing on “global” properties of the population that are robust to the expected differences in the underlying unit responses. Specifically, we ask whether the response-fields estimated in the delayed-saccade task can be used to predict: (1) the overall diversity of unit responses measured in the dots-task, and (2) the resemblance (or difference) of unit responses measured in any two task-configurations and cortical locations. We first develop a description of the population response that is well suited to study these global properties in the measured dots-task responses (Fig. 6), and then use this description to evaluate the response-field based predictions both qualitatively (Fig. 7) and quantitatively (Fig. 8).

**Figure 7.**
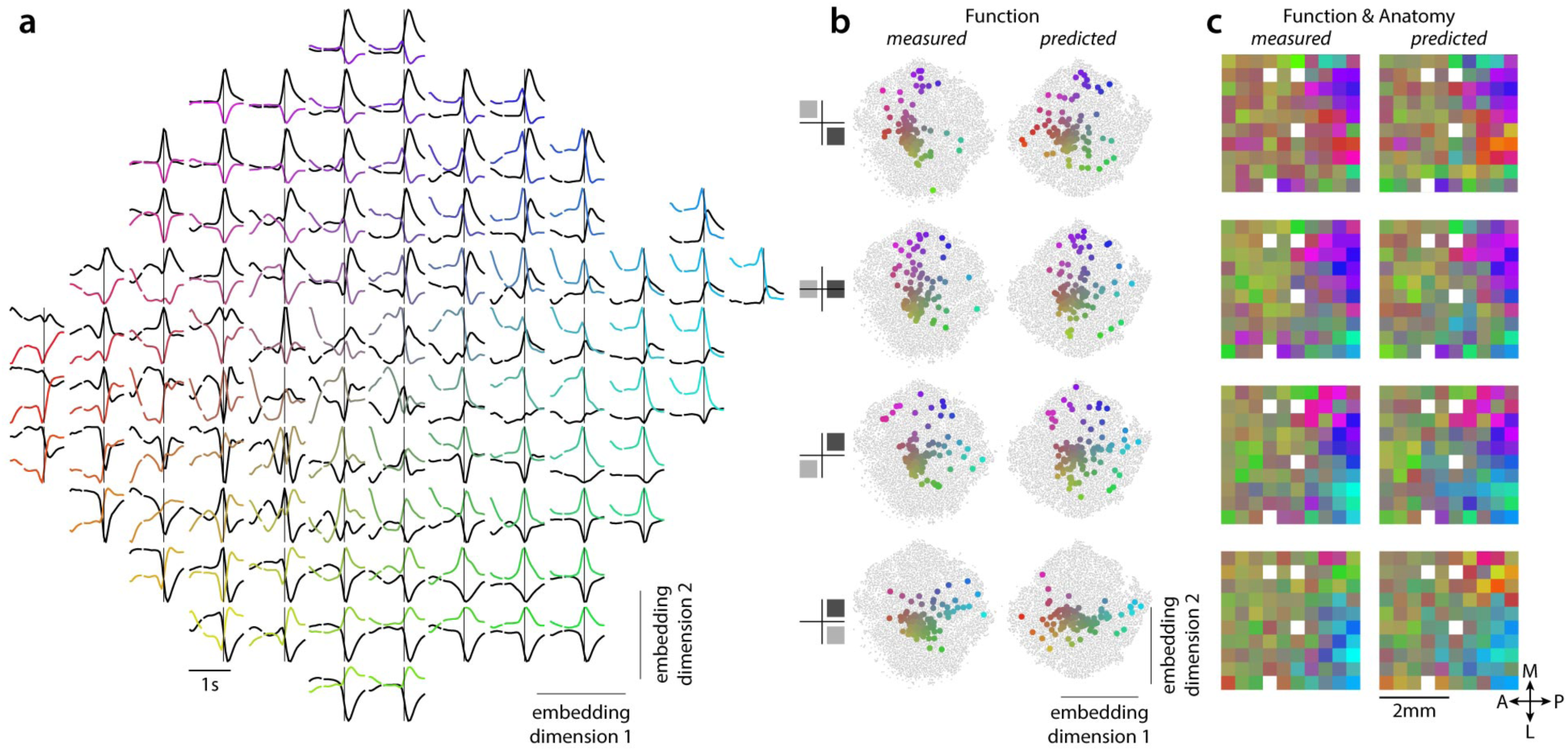
Response-field based predictions of dots-task responses. We used the response-fields estimated in monkey T (Fig. 5) to predict single-unit responses in the dots-task, and embedded the predicted responses with the same approach used for the measured responses (Fig. 6). **a**, Average predicted unit responses, obtained with a non-linear embedding as in Fig. 6d (responses cover a shorter time-range compared to Fig. 6d, scale bars). Only two conditions are shown (choice 1 and 2). **b**, Effect of cortical location and task-configuration on the unit responses observed in the dots-task. Non-linear embedding of measured dots-task responses (gray dots, left; replotted from Fig. 6b) and of predicted dots-task responses (right). Each colored circle corresponds to the average location in embedding space of all units recorded from the same array electrode for a given task configuration (rows). The circles are colored based on their location in the embedding space (same mapping of embedding space to color as in **a** and Fig. 6c). **c**, Measured and predicted topography of unit responses across the cortical surface. Each square corresponds to a circle in **b**, replotted with the same color at the corresponding array location. The resulting mapping of unit responses (colors) to cortical locations depends on task-configuration (rows), and is largely reproduced by the predictions.

**Figure 8.**
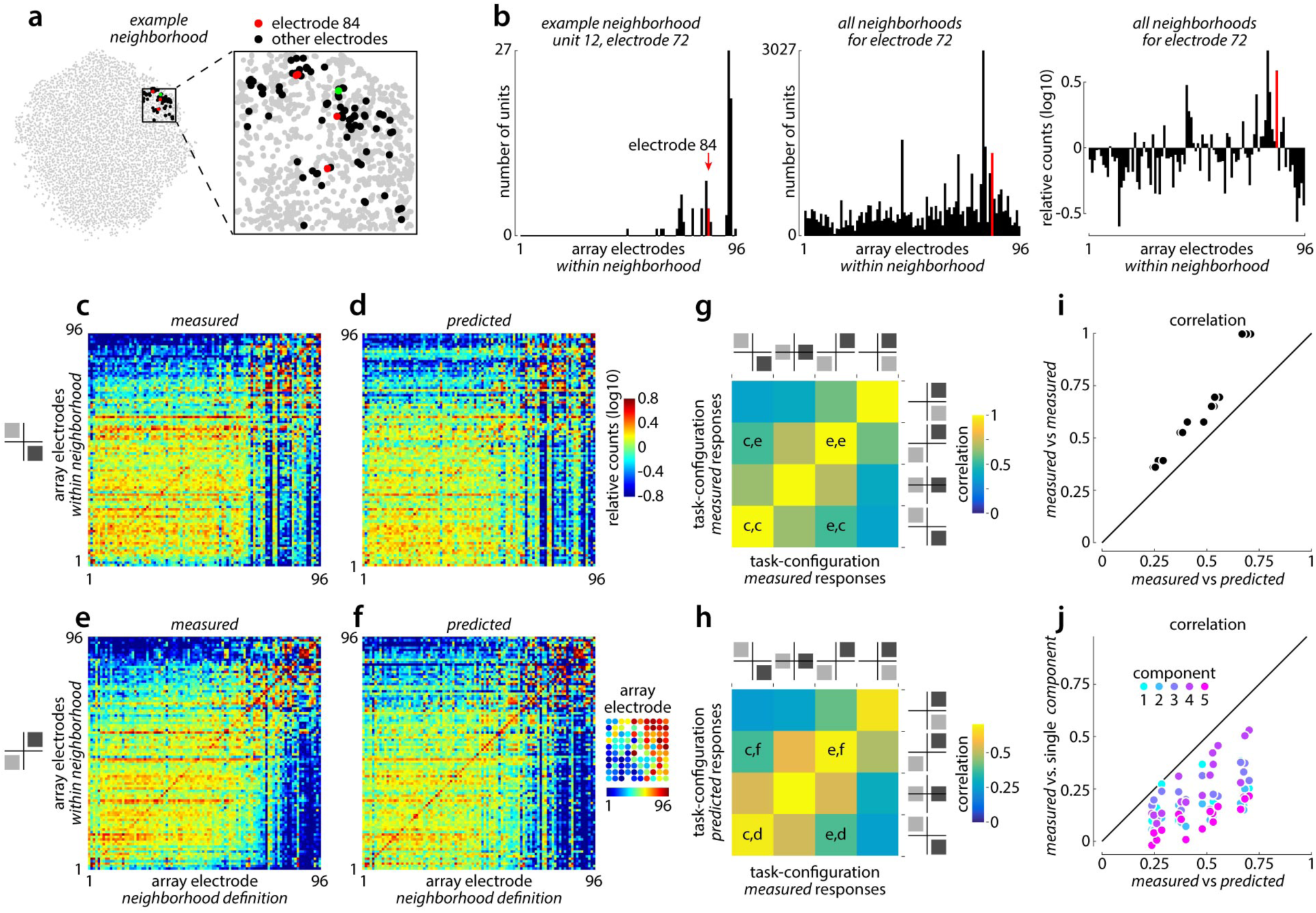
Accuracy of response-field based predictions. **a**, An example neighborhood (red and black dots) shown in the embedding space (gray dots replotted from Fig. 6b). The neighborhood of unit 12 (green dot, from electrode 72; shown in Fig. 1e) is defined as the 90 units that are closest to unit 12 in the high-dimensional space. **b**, Neighborhood-mixing with respect to array electrodes. Left: units in the example neighborhood (**a**) grouped by the electrode at which they were recorded. Middle: same as left panel, but over all neighborhoods of units from electrode 72. Right: mixing between units from electrode 72 and all other electrodes, defined by dividing the corresponding counts (middle panel) by the null hypothesis (no topographical arrangement of unit responses). Large relative counts indicate electrodes with responses that are similar to those on electrode 72, and vice versa. As an example, units from electrode 84 are highlighted in red (as in **a**). **c**, Mixing matrix for measured dots-task responses (monkey T) in one task-configuration (inset, left), with respect to array electrodes. Each column in **c** corresponds to relative counts as in **b** (right panel) for all neighborhoods of the corresponding electrode. Electrodes are ordered as shown in the inset in **f** (see methods). **d**, Same as **c**, for responses predicted based on the response-fields estimated in the delayed-saccade task (Fig. 7). Same task-configuration as in **c**. **e**, Same as **c**, for a different task-configuration. **f**, Same as **d**, for the task-configuration in **e**. **g**, Correlation between measured mixing matrices from different task-configurations. Correlations based on the mixing matrices in **c** and **e** are indicated with corresponding letter pairs. By design, the correlations are symmetric with respect to the positive diagonal. **h**, Correlation between measured (horizontal axis) and predicted (vertical axis) mixing matrices from different task configurations, analogous to **g**. The predictions are most accurate when they are based on the same task-configuration as the measured responses (positive diagonal). **i**, Direct comparison of the correlations in **g** (vertical axis) and **h** (horizontal axis). **j**, The correlations in **h** (horizontal axis), for predictions based on all five choice-components, compared to correlations for predictions based on a single choice component (vertical axis). No single choice-component can account for the accuracy of the predictions for all task-configurations.

### Global structure of unit responses in the dots-task

We exploit the inferred component activations (Fig. 4) to generate a description of the neural population that exploits nearest-neighbor relations between unit responses (Fig. 6; Supp. Fig. 7). As a first step, we reconstruct each unit’s response as a mixture of the 5 component activations (as for generating Fig. 4f,h, rightmost plots). This reconstruction step mainly amounts to aggressively “de-noising” the unit responses (Supp. Fig. 3). Second, we characterize each unit by the resulting average responses for eight distinct conditions (Fig. 6a), differing in choice (choice 1 vs. 2), outcome (correct vs. error), and motion strength (high vs. low). Third, we concatenate these condition-averages to obtain a single vector, which can be interpreted as a single point in a high dimensional space. The dimensionality of this space is given by the product between the number of conditions and the number of time-points per condition. Finally, we use a non-linear dimensionality reduction technique (t-SNE (Van Der Maaten, 2013), see Methods) to find a two-dimensional representation of all points that optimally preserves nearest-neighbor relations (Fig. 6b), meaning that units with similar condition-averaged responses (as assessed by the Euclidian distance between the corresponding points in the high-dimensional space; Fig. 6a) are placed close-by in the two-dimensional representation. We refer to this representation as an *embedding* of the responses, and to the two axes spanning the two-dimensional space as the embedding dimensions.

The embedding reveals the overall diversity of unit responses measured in the dots-task. To show all kinds of dynamics observed in individual units, we averaged the responses of units located nearby in the embedding space (i.e., at the same grid-location in Fig. 6c) and plotted the resulting averages at the corresponding locations (Fig. 6d). Different locations along the embedding dimensions map onto different kinds of responses, including units with persistent predictive activity (top-right locations, choice 1 preferring; left, choice 2 preferring), transient saccade related activity (right and top-left), persistent post-choice activity (top and bottom), and combinations thereof. Notably, our definition of the high-dimensional space in Fig. 6a implies that here unit responses are considered in their entirety, rather than separately in the three task epochs as in Fig. 3. As described above, each unit’s response (Fig. 6a) is reconstructed as a weighted sum of the component activations (Fig. 4a,c). We refer to these weights as the *component contributions*, shown in Fig. 6e. The relative contributions of the five components vary strongly across units, resulting in the diversity of unit responses shown in Fig. 6d.

The embedding also reveals how the unit responses depend on task-configuration (Fig. 6f) and cortical location (Fig. 6g). Some kinds of responses are not observed in some configurations—for instance, responses with large contributions from component 1 (Fig. 6e, left; red points) are rare when the choice 1 target is located in the bottom-right quadrant (Fig. 6f, left), and responses with large negative contributions from component 4 (Fig. 6e, fourth panel; blue points) are rare when both targets are in the right hemi-field (Fig. 6f, right). These observations mimic the properties of the average unit responses for these configurations (Fig. 1d). The responses also vary across cortical locations, i.e. array electrodes (Fig. 6g). Units with large positive or negative contributions from one or more of the choice components (Fig. 6e; red and blue points) mostly occur at posterior locations on the array, close to the arcuate sulcus (Fig. 6g, left). For these posterior locations, the responses also vary based on the medio-lateral position on the array, as units from different medio-lateral locations tend to map onto different locations along the top and right edge of the embedding space (Fig. 6g, right).

### Predicting dots-task responses from delayed-saccade responses

To assess if the response fields estimated in the delayed-saccade task (Fig. 5) can explain the global structure of unit responses in the dots-task (Fig. 6), we generated *predicted* dots-task responses based on the estimated response fields. For any unit recorded in the dots task, we randomly picked a unit recorded in the delayed-saccade task on the same array electrode, and generated predicted responses by interpolating the responses from the delayed-saccade task at the two target locations used in the dots-task. This resulted in a predicted population that matched the one recorded in the dots-task both with respect to total number of units, and to the combinations of recording locations and target configurations.

We evaluate the accuracy of these predictions qualitatively by computing a non-linear embedding of the predicted responses (Fig. 7, Supp. Fig. 8), as we did for the measured responses (Fig. 6). The embedding shows that the response-field based predictions (Fig. 7a) reproduce the overall diversity of measured unit responses in the dots-task (Fig. 6d). In the predictions, the time-course of choice-related activity varies substantially across units (Fig. 7a), and reproduces the different combinations of predictive, saccadic, and post-choice activity observed in the measured responses (compare Fig. 7a with Fig. 6d). As expected, not all features of the dots-task responses are well predicted. Because in the delayed-saccade task only one target is presented on a given trial, and the resulting visual transient differs across choices, for many units the predicted responses separate by choice right after the onset of the target (i.e. at the onset of the response in Fig. 7a; units on the left and right of the embedding space), while in the measured responses the separation emerges gradually during the dots presentation.

Despite these differences, the predictions also qualitatively reproduce how the unit responses depend on task-configuration and cortical location. To illustrate how these two factors interact, we computed the average location in the embedding space of all units recorded from a given array electrode (Fig. 7b; colored circles) and task configuration (7b; different rows). We assigned a unique color to each average location based on the same scheme as in Fig. 6c,d, and then projected that color onto the corresponding electrode location in the array (Kiani et al., 2015) (Fig. 7c; each square corresponds to a dot in Fig. 7b). The resulting images show that the topographical arrangement of the different unit responses (colors in Fig. 7c) is not fixed, but rather depends on task-configuration (Fig. 7c, rows). Critically, this dependency between neural dynamics and task-inputs is well reproduced by the predictions (Fig. 7c, compare measured and predicted).

In addition to the comparisons in Fig. 7c, which rely on two-dimensional embeddings of the data, we also compared the measured and predicted responses directly (Fig. 8; Supp. Fig. 9) based on their original, high-dimensional representations (Fig. 6a). Such a direct comparison is important, because a two-dimensional embedding (Fig. 6b) is necessarily an approximate representation of data that (locally) spans more than two dimensions, as appears to be the case for the measured unit responses. For instance, for any given unit (Fig. 8a, green dot) the 90 closest neighbors in the high-dimensional space (red and black units, defining the “neighborhood” of the green dot) typically do not correspond to the 90 closest units in the embedding space (gray dots closest to the green dot).

To measure the similarity between unit responses recorded on any two electrodes directly in the high-dimensional space, we introduce the concept of “neighborhood-mixing” (Fig. 8a,b). In essence, if unit responses from two electrodes are similar, their corresponding neighborhoods will tend to be overlapping, i.e. mixed. By characterizing the degree of mixing between neighborhoods for any pair of electrodes, we define a “mixing-matrix” (e.g. Fig. 8c) where large counts (relative to the null hypothesis, i.e. no topographical organization of responses) indicate that unit responses on the corresponding electrodes are similar, and vice versa. Comparing mixing-matrices between task-configurations (Fig. 8c,e) and between measured and predicted responses (Fig. 8c,d) is analogous to comparing the rows and columns of Fig. 7c. As in Fig. 7c, the comparisons are robust to the expected differences between measured and predicted responses, but here can be more easily quantified (e.g. in terms of correlations between mixing-matrices, Fig. 8g-j) and do not require a low-dimensional embedding of the data.

The measured mixing-matrices for electrode location vary with task-configuration, as demonstrated by the correlation coefficients in Fig. 8g, in agreement with the structure of the embedded responses (Fig. 7c), and the effects of task-configuration on the mixing matrix are well reproduced by the predictions (Fig. 8h,i). Notably, different predictions generated by reconstructing unit responses with only one of the five choice-components (Fig. 8j) cannot account for the measured mixing-matrices nearly as well, indicating that the mixing-matrices are sensitive to contributions from several choice components (Fig. 4a,b). The accuracy of the predictions thus suggests that global structure of unit responses in the dots-task is largely determined by the properties of the underlying, task-independent response-fields.

## Discussion

Our large-scale recordings reveal a rich and diverse representation of choice at the level of individual units. The recorded units did not fall into distinct functional classes, but rather covered a continuum of response types (e.g., no obvious clusters are apparent in Fig. 6b), with choice-related activity occurring in relation to different task-events in different units, including combinations of early and late choice-predictive activity, saccade-related activity, and post-saccadic activity (Bruce & Goldberg, 1985; Chafee & Goldman-Rakic, 1998; Markowitz, Curtis, & Pesaran, 2015). This diversity of unit responses reflects the combined effect of only a few choice-related components contributing to the population dynamics (Fig. 4). Two predictive components are recruited at consecutive times in the time-window preceding a choice and persist until the initiation of a saccade (Markowitz et al., 2015; Yates et al., 2017). The saccade-related activity is transient, and comprises a fast temporal sequence across units. After the saccade, one or two additional components are recruited and persist until reward delivery. The low-dimensional, persistent dynamics of choice responses before and after the saccade appear more consistent with attractor dynamics (Brody et al., 2003; Ganguli et al., 2008; Machens et al., 2005; Mante et al., 2013; Murray et al., 2017) than with the high-dimensional dynamics predicted by reservoir computing (Buonomano & Maass, 2009; Jaeger & Haas, 2004; Maass et al., 2002; Morcos & Harvey, 2016; Rabinovich et al., 2008).

Even though we do not attempt to assign a functional significance to these choice components, past studies provide pointers in this respect. In particular, the first and second components most likely relate to processes of evidence accumulation or motor preparation (Gold & Shadlen, 2007; Hanks & Summerfield, 2017; Kiani et al., 2014; Schall, 2001; Shadlen & Kiani, 2013). The two components could either reflect two distinct processes (Yates et al., 2017) or, alternatively, both components could reflect a single variable (e.g. integrated evidence) that is represented in a dynamic, time-varying fashion at the level of the population (Goldman, 2009; Harvey et al., 2012; Morcos & Harvey, 2016; Parthasarathy et al., 2017; Spaak, Watanabe, Funahashi, & Stokes, 2017). In either case, it is notable that these choice-predictive components tend to be most active at the time of saccade initiation (Fig. 4a,b), resulting in a peak of activity that precedes the purely movement related, third component. Population responses during saccade initiation thus involve strong modulation of the very same components that are active during saccade preparation (i.e. during the dots and delay periods). This observation seems at odds with findings in premotor and motor cortex during reaches, where movement related activity is strictly orthogonal to preparatory activity (Kaufman et al., 2014). The prominent components observed during the hold period, after the monkey’s choice, seem to represent a form of “postdictive” persistent activity (as opposed to visual activity, see Supplementary Material), which is distinct from the predictive activity encoded by the first two components. Similar postdictive persistent activity has previously been linked to cognitive processes required for decision-making, like updating and maintaining the value of available choice options (Curtis & Lee, 2010).

We find that the interplay between these choice components at the level of individual units is largely preserved between a decision-making task and a visually-guided, delayed-saccade task (Fig. 7,8). This finding is not a forgone conclusion—context-dependence is a hallmark of prefrontal responses, and is thought to be critical for generating learned, cognitively demanding behaviors (Fuster, 2008; Miller & Cohen, 2001; Tanji & Hoshi, 2008). One may thus have expected more prominent differences in the structure of the responses between a decision-making-task, which monkeys learn to master over the course of months, and a much simpler saccade task that is learned over the course of days. A relation between choice-predictive activity in the dots-task and preparatory saccade activity has often been assumed (Kim & Shadlen, 1999; Shadlen & Newsome, 2001). Systematic comparisons of the two, however, have typically not been reported, or suggested little or no relation between these measures (Meister et al., 2013). The observation that the organization of choice responses in the population is preserved across two tasks with different inputs provides a second line of evidence against reservoir computing (Buonomano & Maass, 2009; Jaeger & Haas, 2004; Maass et al., 2002; Morcos & Harvey, 2016; Rabinovich et al., 2008), which predicts strongly input-dependent neural dynamics.

These results seem at odds with the findings of some studies of decision-making in rodents, which have revealed response dynamics that are high-dimensional, organized in temporal sequences spanning the entire trial, and strongly context-dependent (Baeg et al., 2003; Fujisawa et al., 2008; Harvey et al., 2012; Morcos & Harvey, 2016; Rajan et al., 2016; Scott et al., 2017). It is possible that some of these studies may have over-emphasized the prominence of temporal sequences in the population, as they often relied on sorting units based on the time of peak-activation, but did not include a cross-validation step (Fig. 3a,d vs. b,e). Alternatively, the reported dynamics could reflect computational principles that are genuinely different from those implemented by the PFC circuits analyzed here, or could reflect differences in the employed behavioral tasks. For one, we find the strongest evidence for persistent activity during epochs leading to behavioral events of unpredictable timing (the saccade go-cue and the feedback), when it might be advantageous to maintain the neural activity in a stable configuration that is optimal for processing the event information. This constraint does not apply when the timing of relevant task events, and in particular of the choice, is under control of the animal, as is the case for rodents navigating in a real (Baeg et al., 2003; Fujisawa et al., 2008) or virtual environment (Harvey et al., 2012; Morcos & Harvey, 2016; Rajan et al., 2016; Scott et al., 2017). For another, the differences in dynamics could also reflect differences in the dimensionality of the observed tasks (Cowley, Smith, Kohn, & Yu, 2016; Gao & Ganguli, 2015). It seems plausible that tasks involving extended spatial navigation through locomotion (Baeg et al., 2003; Fujisawa et al., 2008; Harvey et al., 2012; Morcos & Harvey, 2016; Rajan et al., 2016; Scott et al., 2017) are higher dimensional than one requiring ballistic saccades to only two locations, and would thus result in dynamics that are much more high-dimensional.

In our recordings, much of the diversity in the responses of individual units in the decision-making task, as well as differences in population dynamics across task-configurations, reflect the topographical arrangement of response-field properties across the cortical surface (Markowitz et al., 2015; Robinson & Fuchs, 1969; Schall, 1997; Suzuki & Azuma, 1983). Thus, even in a prefrontal area like pre-arcuate cortex, whose computations are thought to emerge from the collective activity of large populations of neurons (Mante et al., 2013; Rolls et al., 2008; Wang, 2002), accounts of the dynamics that rely entirely on population-level descriptions may miss relevant structure at the level of individual units. This structure reflects regularities in the underlying anatomical connectivity that are likely to be critical to the functions of the corresponding PFC areas. However, such regularities remain largely hidden in recordings obtained during the dots-task because of the impoverished motor outputs employed (frequently only two saccade targets), a common feature of many tasks currently used in cognitive neuroscience (Hanks & Summerfield, 2017; Shadlen & Kiani, 2013). While these designs have proven extremely valuable in the context of single-unit recordings, the low-dimensionality of the task parameters may lead one to severely underestimate the natural, intrinsic dimensionality of a neural system, even when neural responses are studied with modern, large-scale recording approaches (Cunningham & Yu, 2014).

Current analysis approaches, at the single-unit or population level, can provide insights into different, complementary aspects of such high-dimensional data, but obtaining a complete characterization of neural population responses spanning these levels remains challenging. The non-linear embeddings used here offer a promising approach to study the structure of neural populations in their entirety, while still maintaining an explicit representation of each units’ response. In addition, our nearest-neighbor statistics (Fig. 8a,b) provide a novel and very general approach to building similarity or distance matrices (Kiani et al., 2015; Kriegeskorte et al., 2008) (Fig. 8a-d), which makes essentially no assumptions about the nature of the underlying high-dimensional data. Ultimately, the insights provided even by these novel approaches will be limited by the richness of the employed behavioral tasks, which in many current experimental designs may be insufficient to reveal all the relevant structure in the responses.

## Methods

### Experimental procedures

We collected behavioral and neural data from two adult male rhesus monkeys: monkeys T (14 kg) and V (11 kg). All surgical, behavioral, and animal-care procedures complied with National Institutes of Health guidelines and were approved by the Stanford University Institutional Animal Care and Use Committee. Prior to training on the direction discrimination task, the monkeys were implanted with a stainless-steel head holder (Evarts, 1968) and a scleral search coil for monitoring monocular eye position (Judge, Richmond, & Chu, 1980). We used operant conditioning with liquid rewards to train the monkeys to perform a two-alternative, forced-choice, motion discrimination task, and a visually guided, delayed-saccade task, both described below.

During training and experimental sessions, monkeys sat in a primate chair with their head restrained. Visual stimuli were presented on a cathode ray tube monitor controlled by a VSG graphics card (Cambridge Graphics, UK), at a frame rate of 120Hz, and viewed from a distance of 57 cm. Eye movements were monitored through the scleral eye coils (C-N-C Engineering, Seattle, WA). Behavioral control and data acquisition were managed by a computer running the REX software environment and QNX Software System’s (Ottawa, Canada) real-time operating system.

### Behavioral tasks

Monkeys performed a motion-direction discrimination task in which perceptual judgments were reported by saccadic eye movements to one of two targets (Britten et al., 1992) (Fig. 1a). The eccentricity (6-18 deg of visual angle) and angular location of the targets varied across sessions (Fig. 1b) Animals discriminated the direction of motion in a fixed-duration random-dot kinematogram contained within a circular aperture of 7° (monkey T) or 6° (Monkey V) in diameter and centered on the fixation point. The difficulty of the discrimination was varied parametrically from trial to trial by adjusting the percentage of dots in coherent motion (Britten et al., 1992) (Fig. 1c). The animals were rewarded for indicating the correct direction of motion with a saccadic eye movement to the target corresponding to the prevalent direction of motion (choice 1 or 2). At 0% coherence the animals were rewarded randomly (50% probability).

Each trial (Fig. 1a) began with the appearance of a small spot that the monkey was required to fixate for 500 ms (*fixation period;* ±1.5 deg fixation window) before the two saccade targets were displayed (*target period*). After 400 msec the random-dot stimulus was presented for a fixed duration of 800 msec (*dots period*). The viewing of the dots was followed by a variable-interval *delay period*, during which only the fixation point and the two peripheral targets were visible (300-1100ms, mean 700ms) (Kim & Shadlen, 1999). At the end of the delay period, the fixation point disappeared, cuing the monkey to quickly initiate the choice saccade. The saccade was followed by an additional randomized interval, *the hold period*, during which the monkey was required to fixate the target before the trial outcome (500-1200ms, mean 900ms; ±2-4 deg fixation window, depending on eccentricity). At the end of the hold time, both targets disappeared, a liquid reward was delivered for correct trials, and the monkey was released from behavioral control. At this point, the monkey could initiate the next trial by re-directing his gaze to the central fixation spot, although he did not always do so.

In separate sessions, monkeys also performed a visually-guided, delayed-saccade task (Fig. 5a). The sequence of events in this task was similar to that in the dots task, but no random-dots were shown, and only one target was presented on any given trial. The fixation period (500ms) was followed by the target period (500-1100ms, mean 800ms), the go-cue (fixation point disappearance) and instructed saccade, and the hold period (700-1300ms, mean 1000ms). Within a session, target location on each trial was pseudo-randomly chosen from a set of locations distributed across the visual field. The number of target locations, (24-33), eccentricities (3 values, 4-12 deg) and angular locations (8-11 angles) varied across sessions.

### Neural recordings

We recorded single and multi-unit neural signals with a chronically-implanted 10 by 10 array of electrodes (Cyberkinetics Neurotechnology Systems, Foxborough, MA; now Blackrock Microsystems). The inter-electrode spacing was 0.4 mm; electrodes were 1.5 mm long. Arrays were surgically implanted into the pre-arcuate gyrus (Supp. Fig. 2) according to a previously-published surgical protocol (Santhanam, Ryu, Yu, Afshar, & Shenoy, 2006; Suner, Fellows, Vargas-Irwin, Nakata, & Donoghue, 2005). We targeted the array to a region of prefrontal cortex between the posterior end of the principal sulcus, and the anterior bank of the arcuate sulcus, near the rostral zone of Brodmann’s area 8 (area 8Ar). The arrays were implanted in the left hemisphere in both monkeys. The exact location of the array varied slightly across monkeys (Supp. Fig. 2), due to inter-animal variations in cortical vasculature and sulcal geometry that constrained the location of the array insertion site in each monkey.

Array signals were amplified with respect to a common subdural ground, filtered and digitized using hardware and software from Cyberkinetics. For each of the 96 recording channels, ‘spikes’ from the entire duration of a recording session were sorted and clustered offline, based on a principal component analysis of voltage waveforms, using Plexon Offline Sorter (Plexon Inc., Dallas, Texas). This automated process returned a set of candidate action-potential classifications for each electrode that were subject to additional quality controls, including considerations of waveform shape, waveform reproducibility, inter-spike interval statistics, and the overall firing rate. For clusters returned by this post-processing, both spike-waveform and spike-timing metrics fell within previously-reported ranges for array recordings (Suner et al., 2005).

Daily recordings yielded ∼100-200 single and multi-unit clusters distributed across the array. We do not differentiate between single-unit and multi-unit recordings, referring to both collectively as “units” in a way that is agnostic to their biological origin. No conclusions we draw in this study appear to depend on a distinction between single and multi-unit responses; indeed, we replicated several main findings reported here in a much smaller population of well-discriminated single-units from single-electrode recordings in two additional animals (pre-arcuate and arcuate cortex; data from (Mante et al., 2013), not shown).

Neural responses in the dots-task were recorded over a total of 67 and 62 experiments in monkeys T and V, for a total of 59727 and 37985 trials. Each daily recording was subdivided into several short “sessions” with identical behavioral parameters. The behavioral paradigm was interrupted for a few seconds between the sessions to close and open data files. In many recordings, these interruptions introduced discontinuities in the overall firing rate of units across sessions. To ensure maximal stationarity in the recordings, we thus analyzed each session separately. Overall, we analyzed responses from 185 and 184 sessions in monkeys T and V, yielding a total of 34233 and 44386 units. Since many of these units were likely recorded repeatedly across sessions and days (Chestek et al., 2007; Santhanam et al., 2009), these totals should be interpreted as the number of samples drawn from a smaller underlying neural population of unknown size. The responses during the delayed-saccade task were recorded over a total of 26 and 11 experiments in monkeys T and V, for a total of 23865 and 4768 trials, distributed across 61 and 13 sessions, and yielding 11468 and 3069 units. In both monkeys, recordings from the two tasks were interleaved over the same time-period (9 and 18 months, monkeys V and T).

Two studies reporting analyses on a subset of these recordings were published previously (Kiani et al., 2014; Kiani et al., 2015).

### Analysis of choice behavior

To quantify the effect of target configuration on the monkey’s performance, we computed for each session the percentage of correct responses as a function of motion coherence (Fig. 1c, left) and fitted a sigmoidal curve to all the resulting points from the same target configuration (Fig. 1c, middle; one curve per target configuration). The estimates of percentage correct at zero coherence are more variable than those obtained for non-zero coherences, as the latter are based on twice as many trials per session (average over two directions of motion). We summarized the performance of the monkey in each target configuration by using the fitted curves. Specifically, we defined an average performance for each target configuration (Fig. 1c, right) as the average fitted performance over 100 coherence values spaced logarithmically between 1% and 100% (Fig. 1b).

### Analysis of eye movement data

We estimated the saccade initiation and end times in each trial by applying a Gaussian fit to the eye velocity profile of the choice saccade. Saccade initiation and end times were defined, respectively, as the times when the derivative of the fit first exceeded a velocity threshold of 15°/sec and decreased below 10°/sec.

### Analysis of neurophysiology data

Throughout the paper, we consider neural responses occurring during two distinct, largely non-overlapping time epochs. The first epoch starts 100ms before onset of the random-dots, and ends 200ms after their offset. The second epoch starts 600ms before and ends 600ms after the initiation of the choice saccade. For each trial, we computed time-varying firing rates by counting spikes in non-overlapping, square time-windows of width 50ms.

We defined condition-average responses for each unit by averaging the time-varying firing rates across all trials belonging to a given condition. We define each condition based on a combination of task variables, specifically the monkey’s choice (choice 1 or 2), trial outcome (correct or error), motion coherency (values differ across experiments), and overall difficulty (high vs. low coherence, defined by splitting coherences into two sets). Some of these conditions are shown only in a subset of the figures. In sessions with one target per hemifield, choice 1 was defined as the target in the right visual hemifield, i.e. contralateral to the left hemisphere containing the recording array (see above, Neural recordings); when both targets appeared in the same hemifield, choice 1 and choice 2 corresponded to the targets in the upper and lower visual fields, respectively.

The condition-average responses are defined at the level of individual experimental sessions, each containing only a fraction of the trials recorded on a given day (see Neural recordings above). As a result, the condition-averages can be rather noisy (Supp. Fig. 3, Measured). We thus use a dimensionality reduction approach to de-noise the responses of individual units. For each session, we identify patterns of population activity that are robustly modulated by the choice on a given trial and reconstruct each unit’s response based only on these choice-related patterns—the contribution of all other patterns is removed from each unit. The resulting, reconstructed responses differ from the raw condition-averages mainly in two respects (Supp. Fig. 3; 15 PC dimensions or 10 choice components). First, they are substantially less noisy. Second, they typically display smaller modulations over time that are common to all conditions. This second observation implies that a substantial fraction of the condition-independent variance occurs in a subspace of the population dynamics that is orthogonal to the inferred task-related patterns, in agreement with previous reports (Kobak et al., 2016). As the focus of this report lies on the choice-related components of the response, these condition-independent signals are not considered further.

Reconstructed unit responses based on varying degrees of de-noising are shown in Fig. 1 and Fig. 3 (10 choice components), Fig. 6 (5 choice components), and the corresponding supplementary figures. Including additional dimensions to the unit responses does not significantly affect the results of the sorting procedure used to identify sequences at the level of individual units (Fig. 3 and Supp. Fig. 4). Including additional dimensions to the unit responses does also not affect the conclusions about the structure and origin of diversity in unit responses across the population (Figs. 6-8). However, the non-linear embeddings are sensitive to noise, and for large number of components (e.g. 10) result in a high number of artefactual, small clusters that do not reflect bona fide neural activity (not shown). The details of the dimensionality reduction approach are described in the next sections.

### Targeted dimensionality reduction

We analyzed the population response in each session with *Targeted Dimensionality Reduction*, a dimensionality reduction approach based on linear regression (Mante et al., 2013). We first applied a “soft” z-scoring (referred to simply as “z-scoring” below) to the responses of a given unit by subtracting the mean response from the firing rate at each time and in each trial and by dividing the result by the standard deviation of the responses (plus a constant):

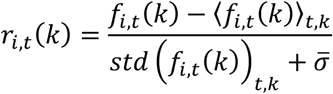

where *f*_*i,t*_(*k*) and *r*_*i,t*_(*k*) are the firing rate and z-scored responses of unit *i* at time *t* and on trial *k*, ⟨.⟩_*t,k*_ and *std*(.)_*t,k*_ indicate the mean and standard deviation across times and trials, and 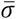 is a constant defined as the median of the standard deviation across all units in a session. The z-scoring de-emphasizes the contribution to the population response of units with very high firing rates (typically multi-unit activity), while the constant term ensures that units with very small firing rates are not over-emphasized. We do not apply any temporal smoothing to the responses.

We used a permutation test to determine the fraction of units with significant choice responses. For each unit, we measure the largest absolute difference between average choice 1 and 2 responses over all times:

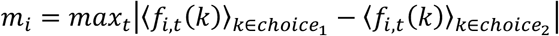

We assessed the significance of *m*_*i*_ by comparing it to the null distribution for *m*_*i*_ obtained with 10,000 random permutations of trials *k*. This test makes no assumptions about the distribution of *f*_*i,t*_(*k*) and incorporates the correction for multiple comparisons across times *t*.

We describe the z-scored responses of unit *i* at time *t* as a linear combination of several task variables:

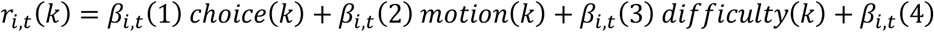

where *choice*(*k*) is the monkey’s choice on trial *k* (+1: to choice 1; −1: to choice 2), *motion*(*k*) is the “signed” motion coherence of the dots on trial *k* (positive values for motion towards the choice 1 target, and negative values towards choice 2), and *difficulty*(*k*) is the “unsigned” motion coherence, i.e. the absolute value of *motion*(*k*). The regression coefficients *β*_*i,t*_(*v*), for *v*=1 to 3, describe how much the trial-by-trial firing rate of unit *i*, at a given time *t* during the trial, depends on the corresponding task variable *v*. The last regression coefficient (*v*=4) captures variance that is independent of the three task variables, and instead results from differences in the responses across time. The signed and unsigned coherence are added to the regression for consistency with a previous study (Mante et al., 2013) but explain only little variance in the responses compared to choice (not shown).

To estimate the regression coefficients *β*_*i,t*_(*v*) we first define, for each unit *i* and time *t*, a matrix ***F***_*i*_ of size *N*_*coef*_ × *N*_*trial*_, where *N*_*coef*_ is the number of regression coefficients to be estimated (4), and *N*_*trial*_ is the number of trials recorded for unit *i*. The first three rows of ***F***_*i*_ each contain the trial-by-trial values of one of the three task variables. The last row consists only of ones, and is needed to estimate *β*_*i,t*_(4). The regression coefficients can then be estimated as:

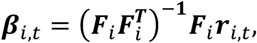

where ***β***_*i,t*_ is a vector of length *N*_*coef*_ with elements *β*_*i,t*_(*v*), *v*=1-4. We denote vectors and matrices with bold letters, and use the same letter (not bold) to refer to the corresponding entries of the vector or matrix, which in this case are indexed by *v*.

The regression coefficients ***β***_*i,t*_ can be re-arranged to produce a set of coefficient vectors ***β***_*v,t*_ (*v*=1-4) of length *N*_*unit*_ whose entries *β*_*v,t*_(*i*) correspond to the regression coefficient for task variable *v* for unit *i* at time *t*. Each vector ***β***_*v,t*_ then corresponds to the direction in state space that accounts for variance in the population response due to the corresponding task variable at time *t*. Targeted dimensionality reduction involves projecting the population response into subspaces derived from these regression vectors.

Single-trial population responses are constructed by re-arranging the z-scored responses into vectors ***s***_*k,t*_, where ***s***_*k,t*_(*i*) = ***r***_*i,t*_(*k*). The dimensionality of the state space corresponds to *N*_*unit*_, the number of units in the population. Condition-averaged responses ***x***_*c,t*_ are obtained by averaging single-trial responses over all trials belonging to condition *c*. We defined conditions based on the choice of the monkey (choice 1 or choice 2), the motion coherence, the outcome of the trial (correct or incorrect), and pairwise combinations thereof.

We used PCA to identify the dimensions in state space that captured the most variance in the condition-averaged population responses. We first build a data matrix ***X*** of size *N*_*unit*_ × (*N*_*condition*_. *T*), whose columns correspond to the z-scored population response vectors ***x***_*c,t*_. *N*_*condition*_ corresponds to the total number of conditions, and *T* to the number of time samples. The PCs of this data matrix are vectors ***v***_*a*_ of length *N*_*unit*_, indexed by *a* from the PC explaining the most variance to the one explaining the least. We use the first *N*_*pca*_ PCs to define a first de-noising matrix ***D*** of size *N*_*unit*_ × *N*_*unit*_:

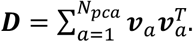

We use this matrix to de-noise the regression vectors defined above by projecting them into the subspace spanned by the first *N*_*pca*_ = 15 principal components:

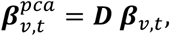

with the set of vectors 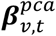 also of length *N*_*unit*_. We make use of the 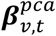 to build the two distinct subspaces of the dynamics that are considered in the main text.

We build a first subspace by focusing on the 10 state-space dimensions that account for most variance in the choice regression vectors. We first define a matrix ***Y*** of size *N*_*unit*_ × 2*T* whose first *T* columns correspond to the de-noised regression vectors of choice 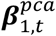 and second *T* columns correspond to 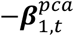, compute principal components ***w***_*a*_ of this matrix, and define a projection matrix ***D***^10^ based on the first 10 PCs:

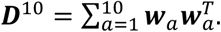

We build a second subspace based on de-noised ‘regression vectors’ obtained by averaging 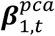 over all times *t* falling within time windows *S*_*j*_ for *j*=1-5:

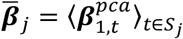

where each 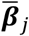 is of dimension *N*_*unit*_, and the time windows *S*_*j*_ cover times *t* within the intervals [0.20, 0.60] relative to dots onset, and [−0.55, −0,30], [−0.10, −0.05], [0, 0.05], [0.10, 0.15] relative to saccade onset (all times in seconds). We orthogonalize the regression vectors with the QR-decomposition:

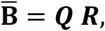

where 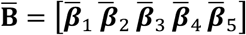 is a matrix whose columns correspond to the regression vectors, ***Q*** is an orthogonal matrix, and ***R*** is an upper triangular matrix. The first five columns of ***Q*** correspond to the orthogonalized regression vectors 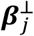. The entries 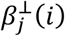 for *j* =1-5 correspond to the “component contributions” in Fig. 6e, and together are analogous to the “component patterns” shown schematically in Fig. 2c. Because of the orthogonalization step, each component pattern explains distinct portions of choice-related variance in the responses. Note that we did not apply any temporal smoothing to the responses, and thus the component activations faithfully reflect the temporal dynamics of the underlying population responses. We define the projection matrix ***D***^5^ based on the orthogonalized regression vectors:

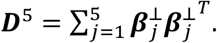

We use the projection matrices ***D***^10^ and ***D***^5^ to de-noise the responses of individual units, and to focus our analyses to the contributions of choice to the responses (see below, Reconstructed unit responses). The first projection matrix, ***D***^10^, results in a “milder” de-noising, and is based on a conservative estimate of the number of choice components in the population response from any single session (Figs. 1,3). This conservative estimate ensures that no “meaningful” diversity of unit responses is lost because of the denoising, which is important in particular for the identification of sequences across units (Fig. 3). The second projection matrix, ***D***^5^, is based on a definition of choice-components that is tailored to identify sequences or recruitment of patterns in the population (Fig. 4), and results in a more aggressive denoising suitable for generating the non-linear embeddings (Fig. 6). The bulk of the variance in the denoised regression coefficients 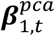 is contained in the subspace defined by ***D***^5^ (Fig. 4i,j; see below, Quality of reconstructions), and the overall diversity of unit responses across the population is similar when responses are de-noised with either projection matrix (Supp. Fig. 4), suggesting that choice related responses within a session can be captured by less than 10 components.

### Component activations

To extract component activations (Fig. 4a,c; schematically in Fig. 2d) we first define a matrix ***X***_*k*_ of dimensions *N*_*unit*_ × *T*, whose columns correspond to the single-trial population responses ***s***_*k,t*_(*i*) (see above, Targeted dimensionality reduction). Component activations are obtained by projecting the single trial responses into subspace defined by the orthogonalized regression vectors:

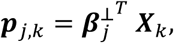

where ***p***_*j,k*_ is a set of time-series vectors over all components and trials, each with length *T*. To avoid the extraction of spurious choice activations, for each experimental session we computed the activations ***p***_*j,k*_ with a 10-fold validation procedure. We first randomly assigned each trial *k* to 1 of 10 sets. We then computed the component activations ***p***_*j,k*_ for all trials in a given set based on orthogonalized regression vectors 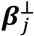 that were estimated from responses on the remaining 9 sets of trials. We repeated this procedure 10 times to compute ***p***_*j,k*_ for all trials in the session.

We quantified the strength of choice related activity along component *j* and at time *t* as the area under the ROC curve between the two distributions of ***p***_*j,k*_ corresponding to choice 1 and choice 2 trials (Fig. 4b,d; Supp. Fig. 5b,d). We computed condition-averaged component activations ***p***_*j,k*_ by averaging ***p***_*j,k*_ over all trials *k* belonging to a given condition *c* (Fig. 4a,c; Supp. Fig. 5a,c).

### Reconstructed unit responses

We obtain reconstructed, de-noised population responses by projecting the population responses into the corresponding subspaces (Supp. Fig. 3). Specifically, the reconstructions based on the mild de-noising (Figs. 1,3) are obtained from:

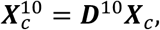

where ***X***_*k*_ is obtained by averaging ***X***_*k*_ over all trials *k* belonging to a given condition *c*. The more aggressively de-noised reconstructions 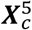 are obtained as a weighted sum of the component patterns, weighed by the corresponding component activations:

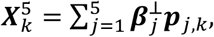

and by again averaging over all trials belonging to a given conditions. The resulting reconstructions are approximately equivalent to those that one would obtain by projecting the condition averaged population responses directly into the subspace spanned by the orthogonalized regression vectors:

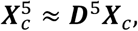

as for the mild de-noising. The equality, however, is only approximate because the ***p***_*j,k*_ are cross-validated (see above, Component activations), while ***X*_*c*_** is not.

### Quality of reconstructions

To validate the quality of the aggressively de-noised reconstructions, we used the reconstructed responses to predict the observed similarity 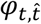 between choice related population activity at times *t* and 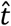 (Fig. 4e,g), defined as:

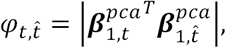

where |.| indicates the absolute value. We applied a cross-validation procedure to compute these similarities, by estimating 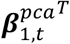 and 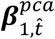 from two separate sets of trials, each containing a randomly chosen half of the trials in a session. We repeated this procedure with 10 different random assignments of trials into two halves, and averaged the resulting similarities to obtain the matrices in Fig. 4e,g. Because of this cross-validation, typically 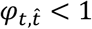 even for 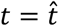. In general, 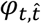 approaches a value of 1 when choice strongly modulates the population response both at times *t* and 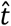, and the resulting choice-related patterns are similar (up to a sign-change). On the other hand, 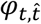 approaches a value of 0 if the patterns at *t* or 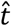 are dissimilar, and/or choice does not strongly modulate the population response at either *t* or 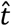.

Predicted similarities can be computed in the same way, by re-applying all the above steps (starting from Targeted dimensionality reduction) to the reconstructed responses. However, this approach results in similarities that are much larger than those in Fig. 4e,g (not shown). These larger values are a consequence of the de-noising procedure, as the estimates of 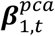 obtained from the reconstructed responses 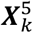 have substantially smaller trial-by-trial variability than those obtained from the original responses ***X***_*k*_. We thus added Gaussian noise to the reconstructions:

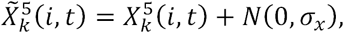

where *N*(0, *σ*_*x*_) are draws from a normal distribution of mean 0 and standard deviation *σ*_*x*_ = 1, and computed predicted similarities from the resulting 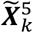, rather than directly from the de-noised reconstructions 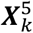. With this choice of *σ*_*x*_, the predicted similarities qualitatively match the observed ones (Fig. 4f,h; rightmost panel). Predicted similarities based on *m* < 5 choice components (Fig. 4f,h; left panels) were obtained in the same way, but with reconstructions based on:

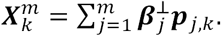

We also directly quantified the fraction of choice related variance in the population responses captured by the individual choice components (colored points, Fig. 4i,j; Supp. Fig. 5i,j), defined as:

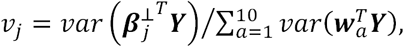

and compared it to the variance explained by individual PCs of the de-noised regression vectors (gray points, Fig. 4i,j; Supp. Fig. 5i,j):

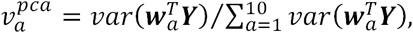

with the relevant quantities defined above (Targeted dimensionality reduction).

### Sequences across units

To demonstrate temporal sequences of activation across individual units, we visualized the population response by plotting all units’ responses after sorting them by the time of peak-activation (Fig. 3; Supp. Fig. 4). For each unit, we first computed the time of peak activation within one of three task epochs, extending from −0.1 to 1s relative to dots-onset, from −0.6 to −0.2s relative to saccade onset, and from - 0.15 to 0.6s relative to saccade onset. We pooled all units from sessions with the same task configuration, and then sorted units based on the computed peak-times, resulting in three different orderings of units (e.g. corresponding to the three panels in Fig. 3a). We applied this analysis to the condition-averaged responses 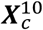 obtained with mild de-noising (see above, Reconstructed unit responses). Here we define 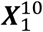 and 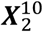 as the average activity over all choice 1 and choice 2 trials, respectively. We computed the peak times and sorted responses either on 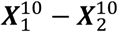 (e.g. Fig. 3a,b), or directly on 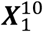 (Fig. 3c, top) and 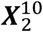 (Fig. 3c, bottom). In all panels, after sorting we averaged the responses of the 250 neighboring rows (i.e. units) with a square moving window, and down-sampled the units by keeping only every 100^th^ row.

To identify contributions of trial-by-trial variability to the resulting plots, we generated each plot from two separate groups of trials, the “validation” set (reflecting only contributions to the responses that are conserved across trials) and the “sorting” set (reflecting also trial-by-trial variability). For each experimental session, we randomly assigned half of the trials to the sorting set, and the other half to the validation set. We used only responses from the sorting set to compute the peak activation times for each unit, and then ordered the unit responses from both the sorting (Fig. 3a) and validation set (Fig. 3b) based on these times. As a result, the ordering of units along the vertical axis across the resulting two plots is preserved, but is entirely determined by responses in the sorting set. We repeated this procedure 10 times for each session, with different random assignments of trials into the sorting and validation sets, and obtained e.g. Fig. 3 by averaging the plots resulting from the 10 different orderings of units.

### Simulations of choice-encoding scenarios

We illustrate how different representations of choice could be revealed by our dimensionality reduction approach with simulated population responses corresponding to four idealized scenarios for the encoding of choice-related activity (Fig. 2). Our goal was not to reproduce the full richness of unit responses observed in prefrontal cortex, but rather to capture the defining features of each scenario with the simplest possible population of idealized neurons. We constructed population responses such that the population average response was (approximately) the same across all encoding scenarios (Fig. 2a, red curves).

For all scenarios, we constructed single unit responses ***u***_*q,e*_ covering *K* =73 temporal samples, for encoding scenarios *e*=1-4. The entries ***u***_*q,e*_(*k*) can be thought of as the average choice 1 response for unit *q*, encoding scenario *e*, at time *k* for choice 1. The average response for choice 2 is set to zero at all times. The various encoding scenarios differ with respect to the definition of ***u***_*q*_. The population average response 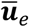 is defined by averaging all the single units responses, i.e.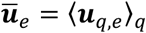.

1. <u>Sequence across units (Fig. 2, second row)</u>. We first define normalized responses *n*_1_(*k*) = *sin*^2^(*π*(*k* – 1)/24) for *k*=1-25 and *n*_1_(*k*) = 0 otherwise. For *q*=2-49 we define *n*_*q*_(*k*) = *n*_1_(*k* – *q* + 1) for *k*=*q*-(*q*+24), and *n*_*q*_(*k*) = 0 otherwise, which corresponds to delaying the response *n*_1_(*k*) by *q*-1 temporal samples. We obtain single unit response as ***u***_*q*,1_ = ***g***^*T*^***n***_*q*_, where *g*(*q*) is a gain factor that emphasizes the responses of units around the time of saccade. We set *g*(*q*) = 1for *q*=1-36 and *g*(*q*) = 1+ *sin*^2^(*π*(*q* – 37)/12) for *q*=37-49.
2. <u>Stable (Fig. 2, first row)</u>. Here the response of every unit in the population is identical up to a scaling factor, and corresponds to the population average response 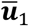 obtained from sequential encoding. Specifically, 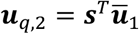 with the scaling *s*(*q*) homogeneously covering the range 0.1 to 1.9.
3. <u>Recruitment (Fig. 2, fourth row)</u>. We first defined component signals ***c***_*j*_ for *j*=1-3 based on the single-unit responses ***u***_*q*,1_ from sequential encoding. Specifically, we define an early component 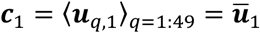, a late component ***c***_2_ = ⟨***u***_*q*,1_⟩_*q=*17:49_, and a saccade component ***c***_3_ =⟨***u***_*q*,1_⟩_q=35:49_. We then defined single unit responses by mixing the early and late signals, ***u***_*q*,3_ = cos(*φ*) ***c***_1_ + sin(*φ*)***c***_2_, for *φ* in the range 0 - *π, q* =1-17; the early and saccade signals, ***u***_*q*,3_ = cos(*φ*) ***c***_1_ + sin(*φ*)***c***_3_, for *φ* in the range 0-2*π, q*=18-49; and the late and saccade signals, ***u***_*q*,3_ = cos(*φ*) ***c***_2_ + sin(*φ*)***c***_3_, for *φ* in the range 0-2*π, q*=50-81.
4. <u>Sequence across patterns (Fig. 2, third row)</u>. We use a different, more elaborate approach to simulate the responses of this encoding scheme. We simulated responses from a non-linear, recurrent neural network consisting of *q* = 100 hidden units and a single read-out (i.e. output) dimension. The input weights of the read-out unit were fixed to 1/100, meaning that the read-out unit computes the average of all hidden unit activities ***u***_*q*,4_ in the network, i.e. the population average response 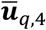. The RNN was randomly initialized (except readout weights) and trained using Hessian free optimization (Martens & Sutskever, 2011) such that after training the activity of its read-out unit matched 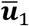. During training, the read-out weights were kept fixed. To regularize and keep RNN dynamics low dimensional, we used Frobenius regularization. Additional RNN parameters and procedures are described in (Mante et al., 2013).

In (1) to (4), we added noise drawn for a normal distribution to each ***u***_*q,e*_(*k*). In analogy to the Targeted Dimensionality Reduction described above, we extract population patterns 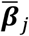 for *j* =1-3, which capture the representation of choice within three different time windows. Here, instead of using linear regression, we simply average the population activity within the temporal windows 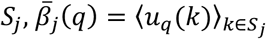 (Fig. 2c, left). The three temporal windows cover the ranges *S*_1_ = [11,15], *S*_2_ = [31,35], *S*_3_ = [51,55]. We then obtain the component patterns 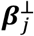 by orthogonalizing these three population patterns (as above; Fig. 2c, right). Component activation are obtained by projecting the single unit activations onto the component patterns, i.e. 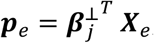, where each row of ***X***_*e*_ corresponds to a vector ***u***_*q,e*_.

### Non-linear embeddings

We used t-SNE (t-Stochastic Neighbor Embedding) to visualize and characterize the diversity of response properties across units in the population. This approach yields a low-dimensional representation of the entire neural population that maintains an explicit representation of each unit’s unit response. Units that are nearby in the low dimensional representation have similar unit responses. We use this representation to characterize the global structure of unit responses in the dots-task (Fig. 6; Supp. Fig. 7), and to compare them to responses predicted based on the response fields estimated in the delayed-saccade task (Figs. 7,8 and Supp. Figs. 8,9; see below, Prediction of dots task responses).

As a first step, we described the response of each individual unit by its condition-averaged responses (Fig. 6a). Here we considered a total of 8 conditions for each unit. Conditions 1-4 correspond to all combinations of choice (1 or 2) and outcome (correct and incorrect). Conditions 5-6 correspond to all combinations of choice (1 or 2) and a reduced measure of motion strength (high and low coherence). To define the latter conditions, for each session we separated trials into high and low coherence conditions based on whether the corresponding unsigned coherence was larger or equal/smaller than the median value across all trials. Here we use condition-averages from 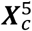, i.e. the aggressively de-noised population responses (see above, Reconstructed unit responses).

As a second step, we use t-SNE to find a two-dimensional representation of the population that optimally preserves nearest neighbor relations (Fig. 6b). To define a distance metric between units, we concatenated each units’ condition average responses into a single vector of length 8 × *T*, i.e. the number of conditions times the number of time samples for each condition (Fig. 6a). We then defined the distance between two units as the Euclidian distance between the corresponding vectors, after the latter had been transformed by a compressive non-linearity. Specifically, we define the distance *d* between units *i*_2_ and *i*_2_ as:

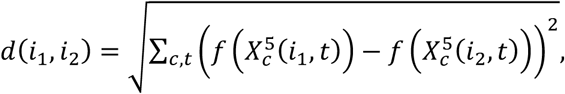

where the sum runs over the 8 conditions *c* = 1to 8, over *t* = 1to *T*, and *f*(.) is the compressive non-linearity:

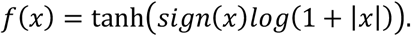

This compressive non-linearity de-emphasizes the contribution of the transient, but large saccade-aligned responses observed in many units (Fig. 1e), and in turn increases the contribution of the persistent activity occurring during the dots, delay, and hold periods to the distance between units. The persistent activity is arguably more likely to depend on the contingencies of the task at hand, as thus provides a more stringent test of the claim that choice-related activity is task-independent. The conclusions in the main text are robust to the exact definition (or the absence) of the compressive non-linearity (not shown).

We used the Barnes-Hut-t-SNE algorithm (Van Der Maaten, 2013) to accommodate the large number of data points (34233 and 44386 in monkeys T and V for the dots task). We used the algorithm with the following choice of parameters (same conventions as in (Van Der Maaten, 2013): perplexity=30; theta=0.5; number of iterations=2500; number of iterations until momentum switch=750). As a pre-processing step, data was projected onto its first 100 principal components, and the algorithm run on the resulting projections. The algorithm is iterative and converges to a local minimum dependent on the initialization of the low dimensional representation. Many of those minima are equivalent as they represent the same neighborhood relationships—for example, a rotation of the entire embedding does not change neighborhood relationships. These invariances complicate visual comparisons of embeddings obtained from different initial conditions. Rather than using random initial conditions, as is typically done, we thus used custom, fixed initial conditions. For both the measured (Fig. 6, Supp. Fig. 7) and predicted (Fig. 7,8, Supp. Fig. 8,9) dots-task responses, we initialized each unit’s position in the low dimensional embedding space as the contributions from the first two choice components (Fig. 6e, first two panels), i.e. the orthogonalized regression vectors 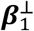 (initialization for first embedding dimension) and 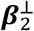 (second dimension). These custom initial conditions do not decrease embedding quality compared to random initial conditions (not shown).

### Characterization of response-fields

We characterized the response-field of each unit with a visually-guided, delayed-saccade task (Fig. 5 and Supp. Fig. 6; see above, Behavioral tasks). In this task, monkeys made delayed saccades to a single target chosen from a set of locations covering the entire visual field (e.g. Fig. 5a). The targets were shown at one of three eccentricities (4°, 8°, 12° degrees of visual angle), but their angular location, and the total number of targets, differed somewhat across sessions. As a result, the average responses to each target location could not be directly compared across all sessions. To obtain a representation of the response-field that is independent of the exact target locations, for each session we interpolated the measured average responses along the vertices of a regular grid (linear interpolation), covering 11 vertices ranging from - 14° to +14° along the horizontal and vertical meridians. At eccentricities larger than 14° the receptive field values were set to zero. We performed the interpolation separately at each time during the trial. We considered responses in two task epochs, covering the range of 0.1 to 0.4s aligned to target onset and −0.5 to 0.5s aligned to saccade onset.

We estimated the peak location of the response-field at any given time by fitting a descriptive function to the condition-averaged responses (Bruce & Goldberg, 1985), expressed as a function of the angular and radial visual field location, *ϑ* and *ρ*:

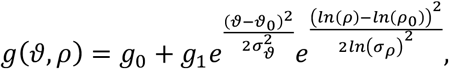

where *ϑ*_0_ and *ρ*_0_ are the angular and radial locations of the peak (red points in Fig. 5c; circles in Fig. 5e-h), *σ*_*ϑ*_ and *σ*_*ρ*_ determine the tuning widths along the angular and radial directions, and *g*_0_ and *g*_1_ set the baseline response and modulation depth of the response-field. We fitted the parameters of this model separately for each unit to average responses within the epochs 0.1-04s after target onset, and 0.05-0.10s after saccade initiation. The model was fit by minimizing the summed square error over all target locations between the model predictions and the condition-averaged responses.

In addition to the response-field based on saccades starting from the fixation point (Fig. 5), for each unit we also estimated a second response-field based on the first saccade leaving the target after the end of trial (not shown). Only the direction and amplitude of the saccade was used to estimate the second response-field, while the starting point of the saccade (one of the target locations) was discarded. For the overwhelming majority of units, the spatial and temporal structure of these two response-fields were very similar around the time of the saccade (−0.4 to 0.3s with respect to saccade onset). In particular, the spatial tuning of the postdictive activity was similar in the two response-fields, even though the retinal images caused by saccades with the same amplitude and angle could be very different depending on their starting point (i.e. the fixation point for the first response-field; one of the targets for the second response-field). Thus, postdictive activity is most likely not driven by visual, retinal inputs, but rather seems more akin to the persistent activity observed before the saccade.

### Prediction of dots-task responses

To explain the diversity of single unit responses observed in the dots task, and its relation to task configuration and cortical location (Fig. 6), we tested whether the global structure of dots task responses can be predicted by the response-field properties measured with the delayed-saccade task. In general, we could not with certainty identify units whose responses were recorded both in the dots task and in the delayed-saccade task (see Neural recordings). Rather than attempting to predict responses on a unit-by-unit basis, we thus tried to predict the overall structure and diversity of the population response during the dots task. For every unit recorded in the dots task, we first randomly picked a unit recorded on the same array electrode in the delayed saccade task. We obtained surrogate dots-task responses as the condition-averaged responses recorded during the delayed-saccade responses at the two target locations used in the dots task. With this approach, we obtained two “predicted” condition averaged responses in the dots task—choice 1 responses from delayed-saccade responses at the corresponding target location, and choice 2 responses at the second target location. In cases where one or both target locations in the dots-task did not have an exact match in the delayed-saccade task, we generated the predicted dots-task response by linear interpolation of the delayed-saccade responses at the corresponding target locations. The result of this procedure is a population of surrogate responses that is exactly matched to the recorded dots task response with respect to the total number of units, and their distribution across recording locations and task configurations.

These surrogate responses lack several potentially important properties of the dots task. First, the surrogate responses are based on recordings where no dots stimulus was present on the screen, and the animal therefore was not involved in deciding between two options. The appearance of the dots stimulus, and the following decision process, result in a characteristic time course of choice-predictive activity in the recorded areas (Fig. 6d). Unsurprisingly, this time-course is not fully replicated in the surrogate responses (Fig. 7a). Second, the surrogate responses are based on recordings where only one target was present on the screen, whereas two targets were simultaneously shown in the dots task. The simultaneous appearance of more than one target may result in competitive interactions within the population of recorded neurons, which again would not be reproduced in the surrogate responses. Third, the timing of the early surrogate response is somewhat mismatched to those recorded in the dots task. For the recorded dots task data, we used responses in the interval between −0.1 and 1s around the *dots-stimulus* onset. For the surrogate data, we instead considered responses in the interval between 0.1 and 0.4s around the *target* onset. The early choice predictive activity in the surrogate responses thus reflects the onset of the single target in the delayed saccade target, while in the dots task it reflects the earliest phases of the decision process.

Despite these differences, the surrogate data reproduces the structure of the recorded dots task responses very well (Figs. 7,8 and Supp. Figs. 8,9).

### Mixing-matrices based on neighborhood relations

To quantitatively compare the predicted and measured dots-task responses at the level of the entire population, we developed a novel non-parametric, statistical approach to characterize the structure of high-dimensional data sets in a way that allows easy comparisons between data sets (Fig. 8). Our approach is based entirely on a quantification of the nearest neighbor relations in the data (Fig. 8a-b). Because high-dimensional data often lie on non-linear manifolds that can locally be approximated by linear manifolds, nearest-neighbor relations are typically easier to define, and can be estimated more robustly, than relations between distant (i.e. very dissimilar) points in the data. Despite being based on *local* relationships between data points, our approach leads to a robust characterization of the *global* structure of a given data set that (unlike t-SNE, e.g. Fig. 6b) does not involve a dimensionality reduction step.

Our approach can be applied to any high-dimensional set of labeled data points. Here, each data point consists of the condition averaged dots-task responses (either measured or predicted) for choice 1 and choice 2. The dimensionality of each data point thus corresponds to twice the number of time-samples in each condition average. Neighborhoods in this high-D space correspond to groups of units with very similar condition-averaged responses. The labels instead correspond to additional properties that can be specified for each data point. Here, we focus on a single label, the electrode location where the unit was recorded (a number between 1 and 96). We then summarize the structure of the data in the high-D space by quantifying which labels co-occur together in local neighborhoods. We call the result of this analysis a “mixing-matrix”, quantifying to what extent units with two specific values for a given label (e.g. recording location, Fig. 8c,e) are locally mixed in the high-D space (i.e. tend to have similar condition-averaged responses). Here, we compute mixing-matrices separately for measured and predicted dots-task responses (Fig. 8c,e measured; Fig.8d,f: predicted) and for each task-configuration (Fig. 8c,d vs. Fig. 8e,f, see insets on the left).

Concretely, we first define a unit’s neighborhood, consisting of its 100 nearest neighbors based on the Euclidean distance in the high-D space. For example, the red and black dots in Fig. 8a correspond to the neighborhood of an example unit from electrode 72, indicated with the green dot. Second, we count how often each label occurs in the neighborhood of a given unit. For each unit, this results in a histogram of labels over all units in its neighborhood. For example, the left panel in Fig. 8b shows the histogram of electrode locations in the neighborhood of the example unit from electrode 72. Third, we pool such histograms for all neighborhoods of units with the same label. For example, the middle panel in Fig. 8b shows the histogram of electrode locations across *all* neighborhoods of units from electrode 72. Fourth, we rescale the resulting pooled histogram by dividing all its values by the predictions of a null hypothesis (Fig. 8b, right), which assumes that the composition of any neighborhood is independent of the properties of the unit that was used to define it (permutation null hypothesis). Based on this null hypothesis, the expected histogram value for units from electrode *i* in the neighborhood of units from electrode *j* is:

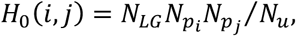

where *N*_*LG*_ is the size of the neighborhood (here *N*_*LG*_ = 100), *N*_*pi*_ is the total number of units in the data set recorded from electrode *i*, 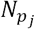 is the total number of units in the data set recorded from electrode *j*, and *N*_*u*_ is the total number of units in the data set. The resulting rescaled histogram corresponds to column 72 of the mixing matrix (e.g. Fig. 8c), where 72 is the electrode location used to define the neighborhoods contributing to the pooled histogram. We then obtain additional columns of the mixing matrix by repeating this procedure for the pooled neighborhoods of units from electrode 1, 2, and so on for all other array electrodes.

The values of the rescaled mixing-matrix (Fig. 8c) at (*i, j*) provide a quantitative measure of the overall (macroscopic) similarity of condition-averaged responses of units from electrode *i* and units from electrode *j*. These mixing matrices can easily be compared between data sets (e.g. measured vs. predicted, or between different task configurations), for example by computing the correlation coefficient between all the values in a pair of mixing matrices (e.g. Fig. 8g).

To ease visual comparison between these mixing matrices for different task-configurations (e.g. Fig 8c,e), we ordered the electrodes along the vertical and horizontal axes in Fig. 8c-e such that electrode locations that recorded units with similar responses are placed nearby. We obtained such an ordering by (1) computing the rescaled mixing matrix values *M*(*i, j*) from measured responses obtained by pooling units from all task-configurations; (2) defining the dissimilarity between electrodes *i* and *j* as: 2 – *log*[*M*(*i, j*)]; and (3) applying multi-dimensional scaling to obtain a one-dimensional ordering of electrodes based on this dissimilarity. The resulting ordering is shown by the coloring of the electrodes in the inset on the right of Fig. 8f.

Predictions based on a single choice component (Fig. 8j) were computed analogously to 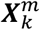 (See Quality of predictions) but by including only a single component activation.

## Supplementary Figures

**Supplementary Figure 1.**
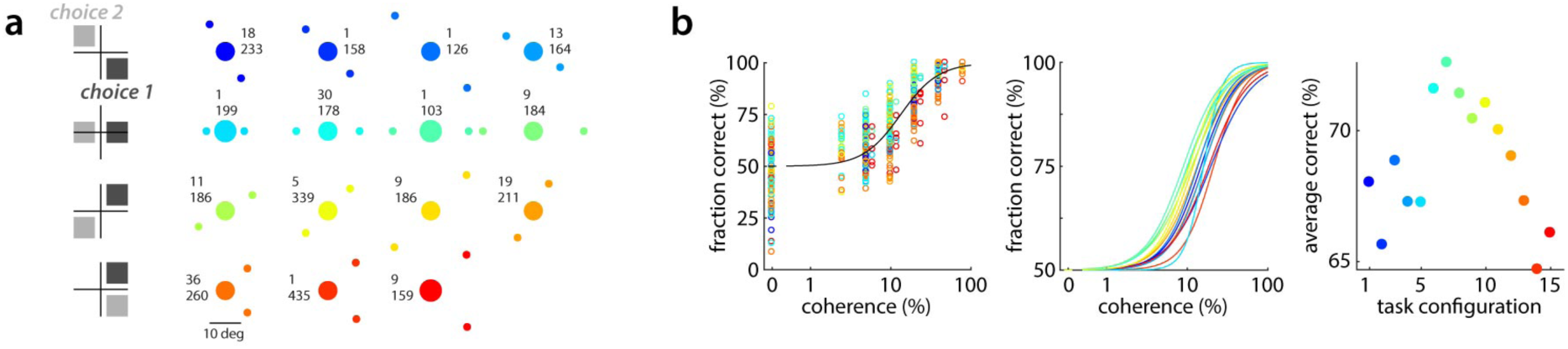
Target configurations and behavioral performance in monkey V. Same conventions as in Fig. 1. **a**, Target configurations (insets: number of sessions, top; average number of behavioral trials per session, bottom), sorted into 4 “task-configurations” (rows). **b**, Behavioral performance, same colors as in **a**. Left panel: fraction correct as a function of motion strength (coherence) and configuration. Middle: Fits of a behavioral model for each configuration, based on the data in the left panel. Right: average performance for each configuration, as estimated from the fits (middle) over a set of coherences common to all configurations.

**Supplementary Figure 2.**
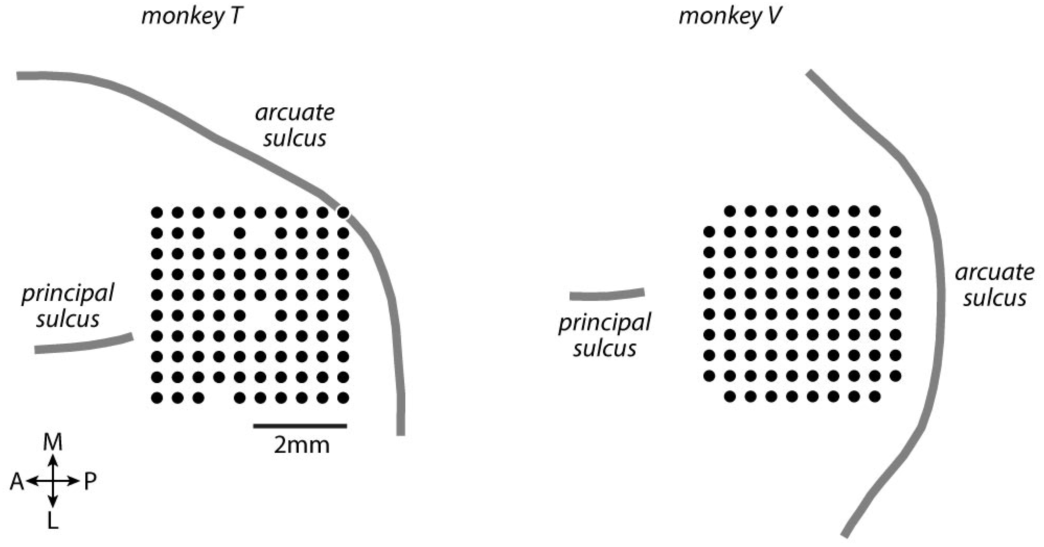
Recording locations in prefrontal cortex. In both monkeys, we obtained single-unit and multi-unit recordings from a 10×10 array implanted in pre-arcuate cortex. Black circles indicate the cortical locations of the 96 electrodes used for recordings.

**Supplementary Figure 3.**
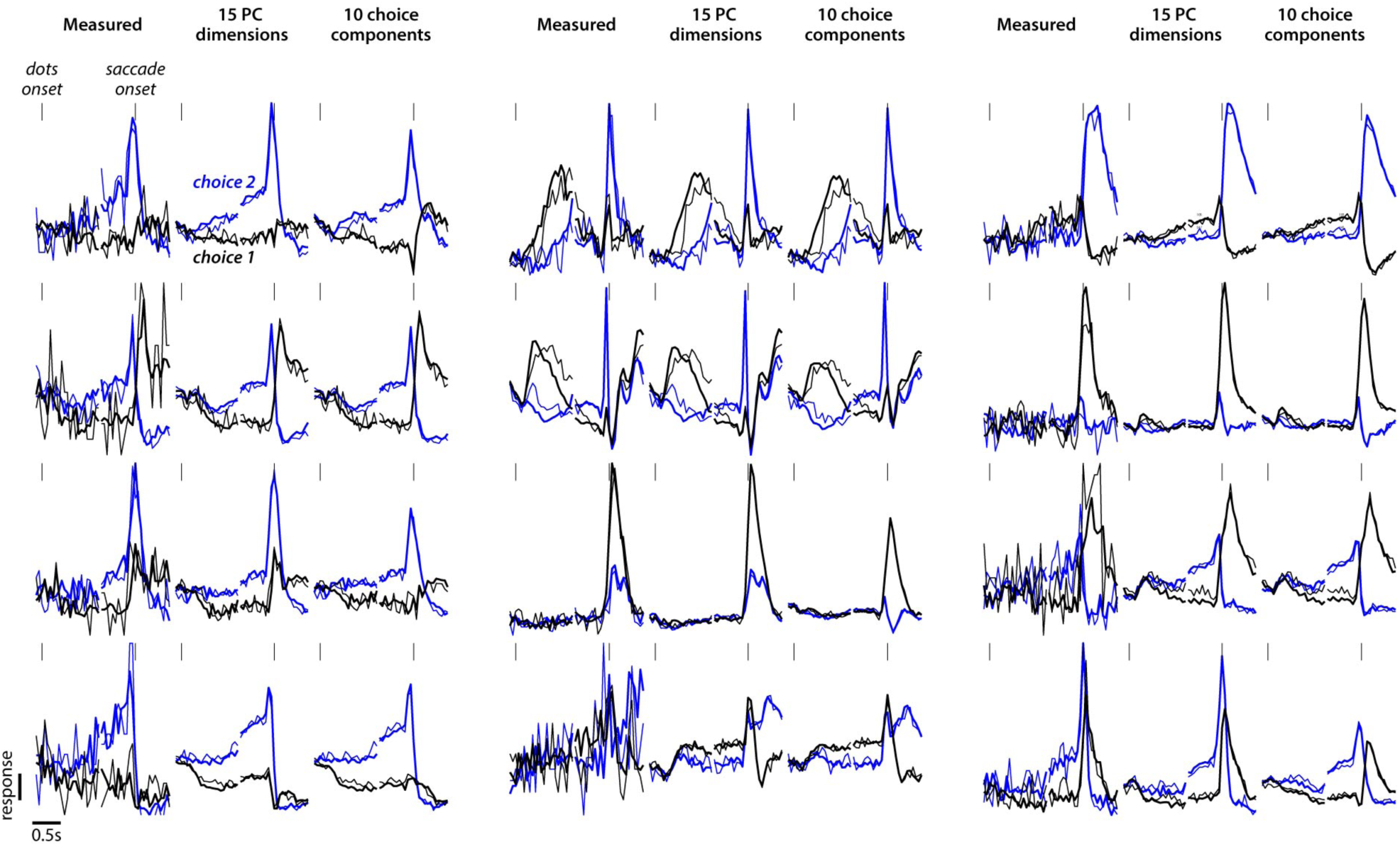
De-noising of single unit responses. Same 12 units as in Fig. 1e. Each group of three adjacent panels shows the raw, measured responses of a given unit (left panel), the responses reconstructed from 15 principle components (middle), and the responses reconstructed from 10 choice components (right, replotted from Fig. 1e with different colors). For any given recording session, we used the raw responses of all neurons to extract dominant components of the population response, either with principle component analysis (PC components) or with Targeted Dimensionality Reduction (choice components). The PC components maximally account for overall variance in the responses, while the choice components only explain variance due to choice. Unit responses are reconstructed based on either the first 15 PC components, or the first 10 choice components. In both cases, all remaining components are assumed to be dominated by noise, and are ignored. This procedure de-noises the single-unit responses, and in some units also removes components of the responses that are common to all conditions (reconstruction from choice components; right panels). Responses de-noised with 10 choice components are used for the analyses in Figs. 1,3. Responses de-noised with only 5 choice components (not shown here) are used for the analyses in Figs. 6, 7, and 8.

**Supplementary Figure 4.**
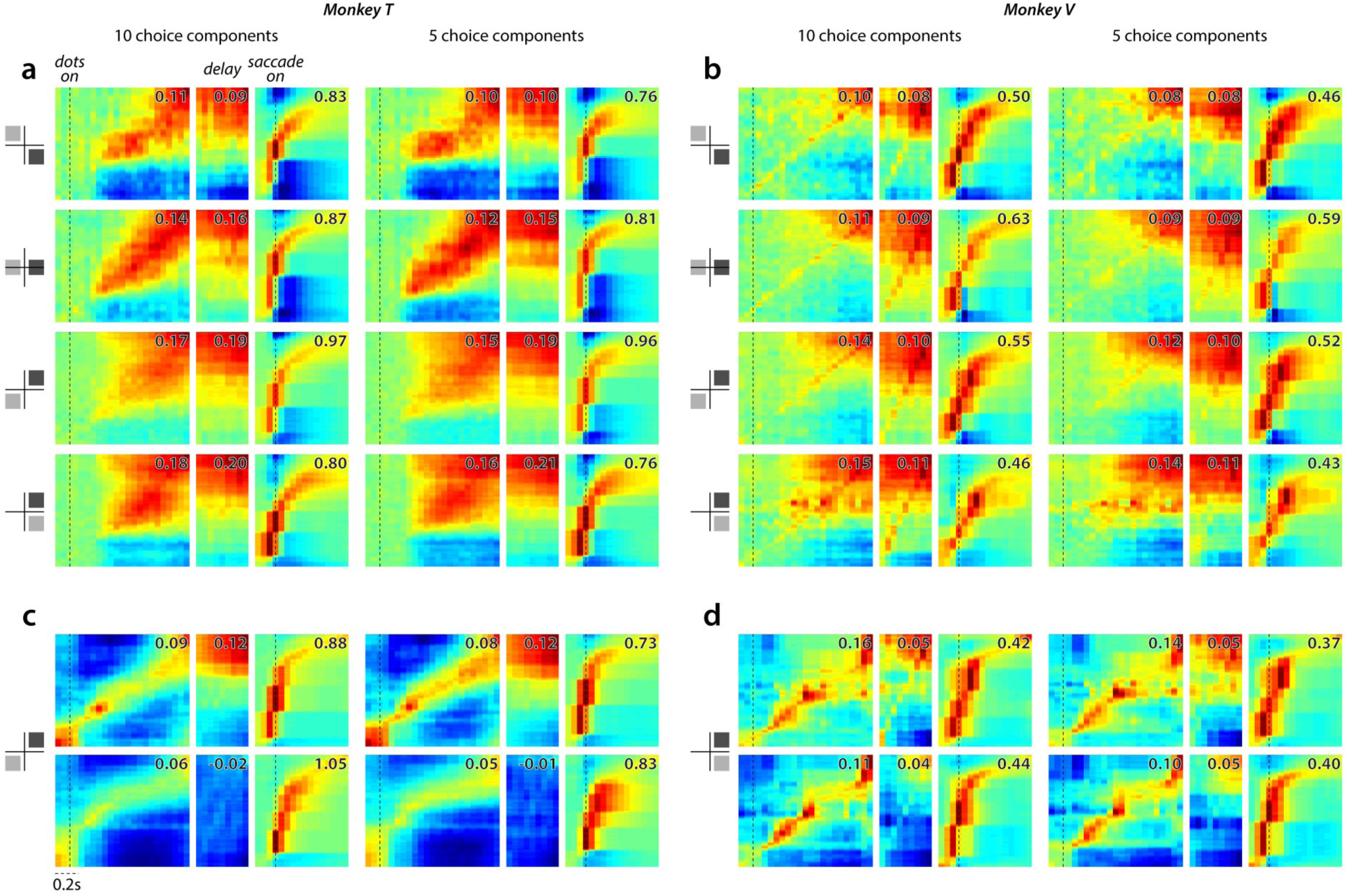
Dynamics of unit responses in PFC for different task-configurations and levels of de-noising. Same conventions as in Fig. 3. **a**-**d**, Response sequences are estimated from unit-responses that were de-noised based on the first 10 choice components in the population of a given session (left columns in **a**-**d**; right panels in Supp. Fig. 3) or based on only the first 5 choice components (right columns). The resulting sequences are similar in both columns, suggesting that choice-related activity in a given session is largely captured by the first 5 choice components of the population response (see also Fig. 4 and Supp. Fig. 5). **a**, Choice-related activity in monkey T, defined as the difference between average responses to choice 1 and choice 2. Averages are obtained based on a randomly selected half of the trials (validation set), and ordered along the vertical axis based on the peak times of choice related activity in the other half of trials (sorting set as in Fig. 3a, not shown here). Analogous to Fig. 3b. **b**, Same as **a**, but for monkey V **c**, Average validation set responses in monkey T for choice 1 trials (top) and choice 2 trials (bottom), sorted based on the peak times estimated on sorting set responses. Analogous to Fig. 3c. Only one task-configuration shown. **d**, Same as **c**, but for monkey V.

**Supplementary Figure 5.**
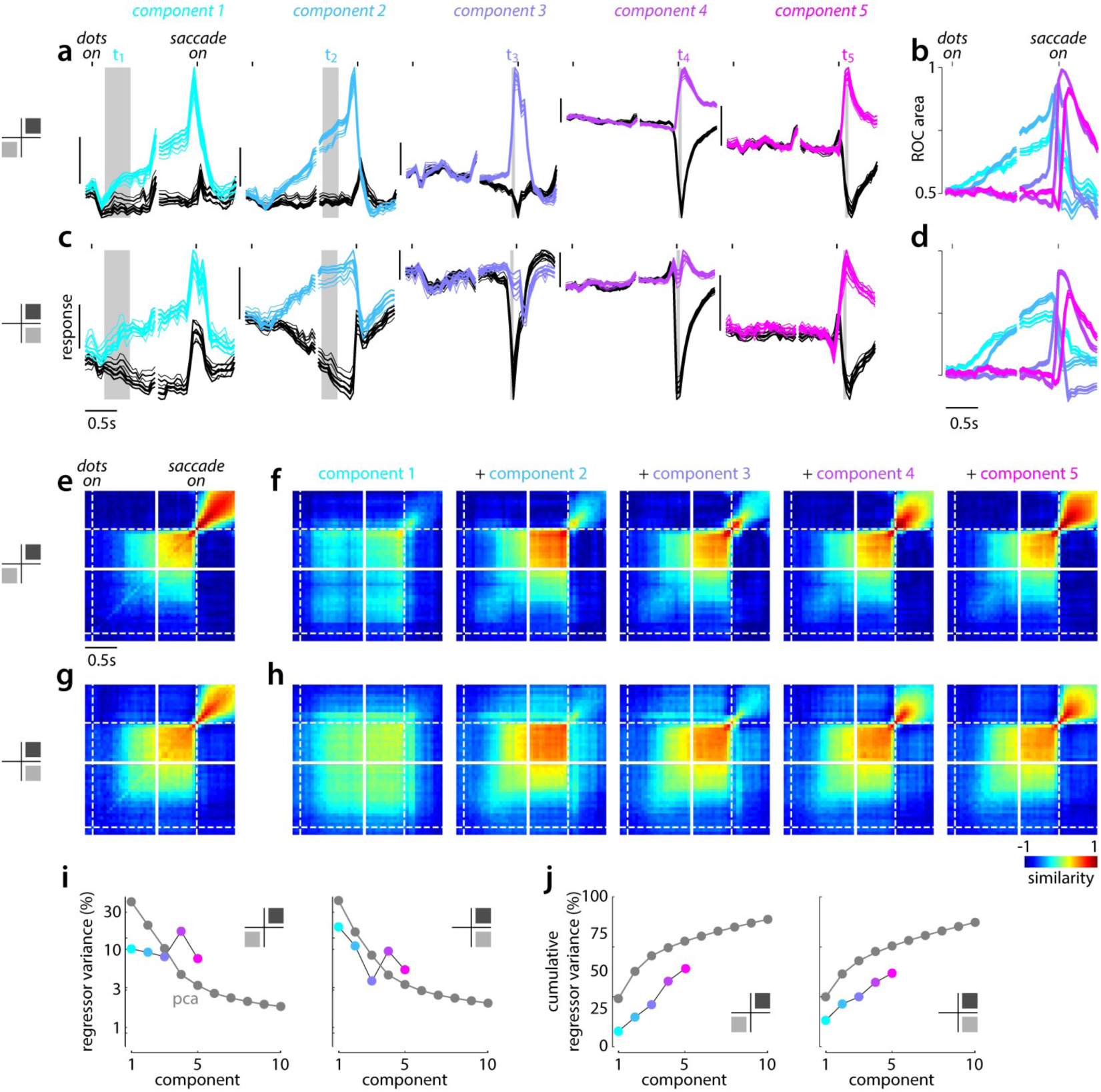
Dynamics of population activity patterns in PFC of monkey V. Same conventions as in Fig. 4, but for data from monkey V. **a**, Component activations for choice 1 (colored) and choice 2 (black) on correct (thick) and incorrect (thin) trials. Error bars indicate standard error of the mean across sessions (analogous to Fig. 2d). **b**, Separation between choice 1 and choice 2 component activations, measured as area under the ROC curve for the corresponding distributions of trial-by-trial activations. **c**,**d**, Same as **a**,**b**, for a different task configuration (left inset, as in Supp. Fig. 1a). **e**, Measured similarity between choice-related population patterns at different times during the trial (analogous to Fig. 2e). **f**, Predicted similarity, based on population responses reconstructed with increasing numbers of choice-related component patterns (from component 1 only, to all five components; left to right). **g**,**h**, Same as **e**,**f**, for different task configurations (left inset, as in Supp. Fig. 1a). **i**, Variance in the choice-related population patterns explained by the 5 component patterns (colored) or by 10 principal components. Task configurations as in **a** (left) and **b** (right). **j**, Cumulative variance explained, computed from the plots in **i**. In **i** and **j**, standard error of the mean over sessions is smaller than the symbols.

**Supplementary Figure 6.**
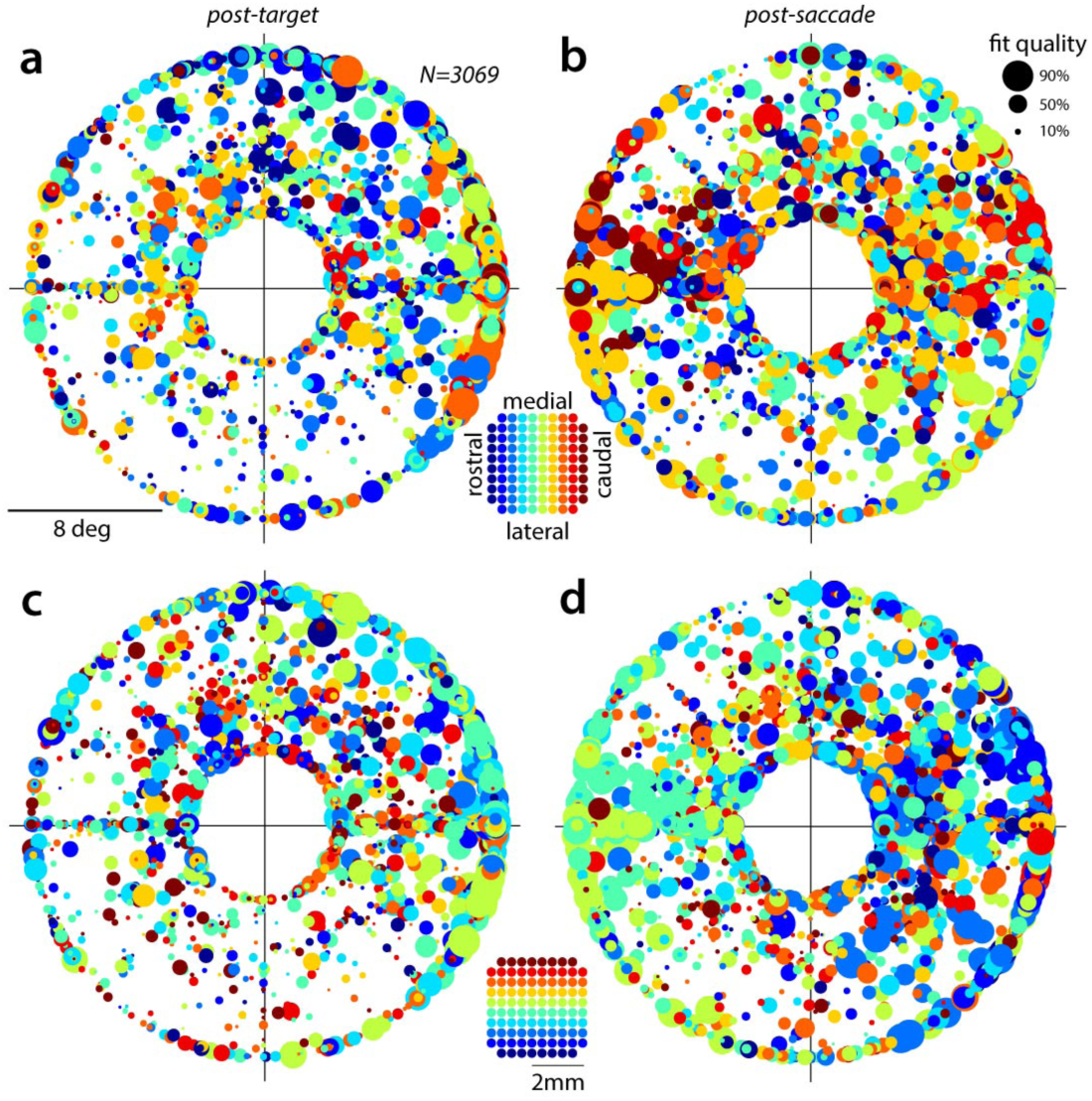
Response-field properties for monkey V. Same conventions as in Fig. 5e-h, but for data from monkey V. **a**, Response-field centers after target onset (0.1-0.4s) for all units recorded in the delayed-saccade task (circles). Circle radius is proportional to the quality of the underlying fit (percentage of variance explained, legend) and can be interpreted as the strength of spatial tuning (strong vs. weak; large vs. small circles). Units are colored based on their anterior-posterior location along the electrode array (inset) and were plotted in random order. **b**, same as **a**, for response during the hold period (0.05-0.1s after saccade onset). **c**-**d**, same as **a**-**b**, but units are colored based on their medio-lateral location on the array (inset). The strongest target related responses occur in the upper right quadrant (**a**), as in monkey T (Fig. 5e). After the saccade, responses also occur at ipsilateral target locations. The location of response-field centers after the saccade (i.e. ipsilateral or contralateral) varies along the medio-lateral axis of the array (**d**), as in monkey T (colors, Fig. 5h).

**Supplementary Figure 7.**
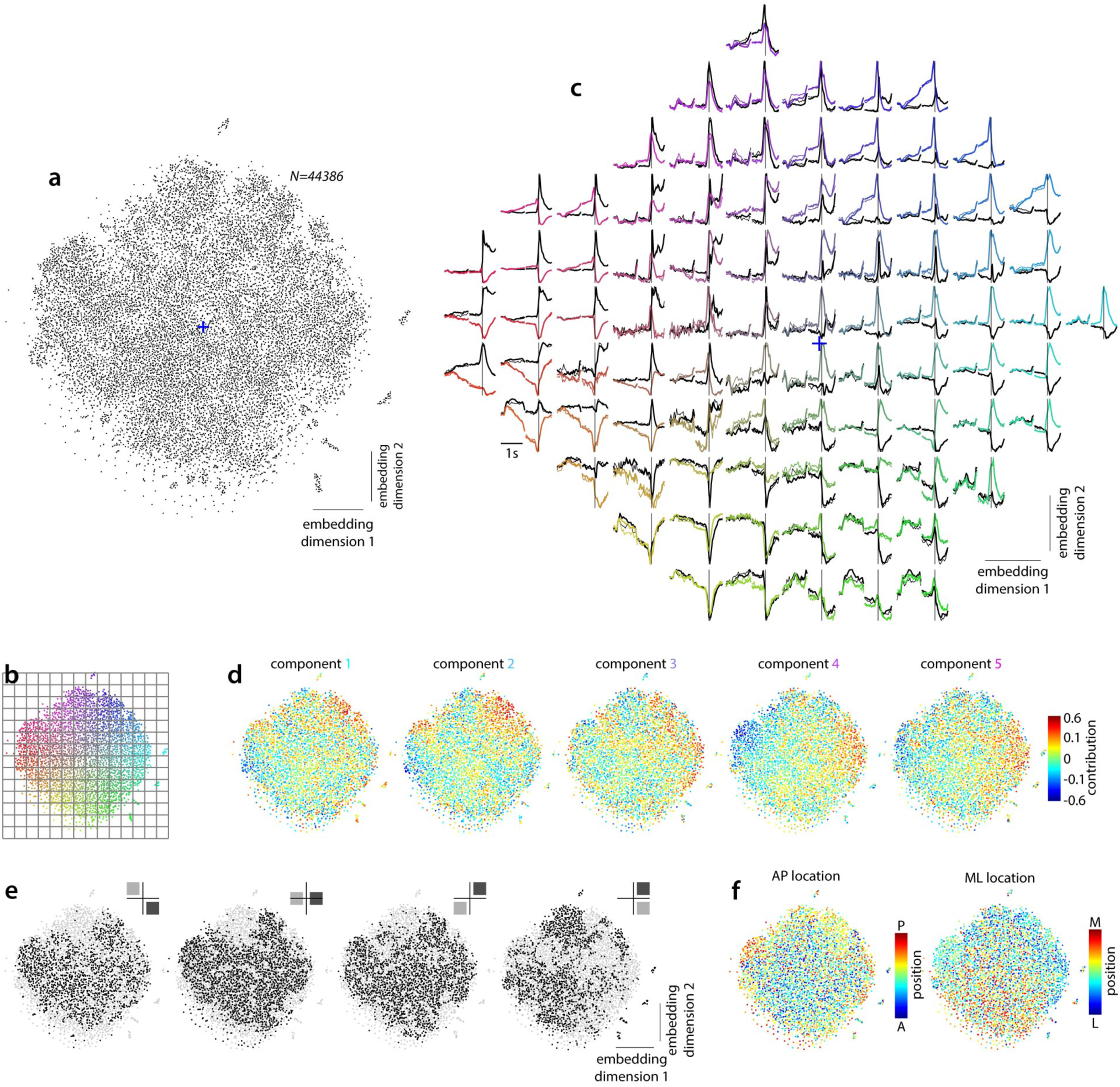
Global structure of unit responses in monkey V. Same conventions as in Fig. 6, but for data from monkey V. **a**, Two-dimensional, non-linear embedding of all units in the population obtained with t-SNE. **b**,**c**, Average unit responses for nearby units at different grid-locations (panel **b**) along the embedding dimensions. Choice 1 averages (panel **c**) are colored based on the radial and angular position of the corresponding units in the embedding space (panel **b**). The blue cross marks the origin of the embedding space, as in **a**. **d-e**, Task-related and neural factors contributing to the diversity of unit responses. The arrangement of dots (i.e. units) in each panel corresponds is identical to the non-linear embedding in **a**. Units in **d** and **f** are plotted sequentially in random order. **d**, The contributions of the five component activations to the responses of each unit in the population. **e**, Embedding location of units recorded with a given task configurations (black points; task-configurations as in Supp. Fig. 1a) overlaid over all units in the population (gray points). **f**, Cortical location of all the units in the population. AP, anterior-posterior axis along the electrode array; ML: medio-lateral axis.

**Supplementary Figure 8.**
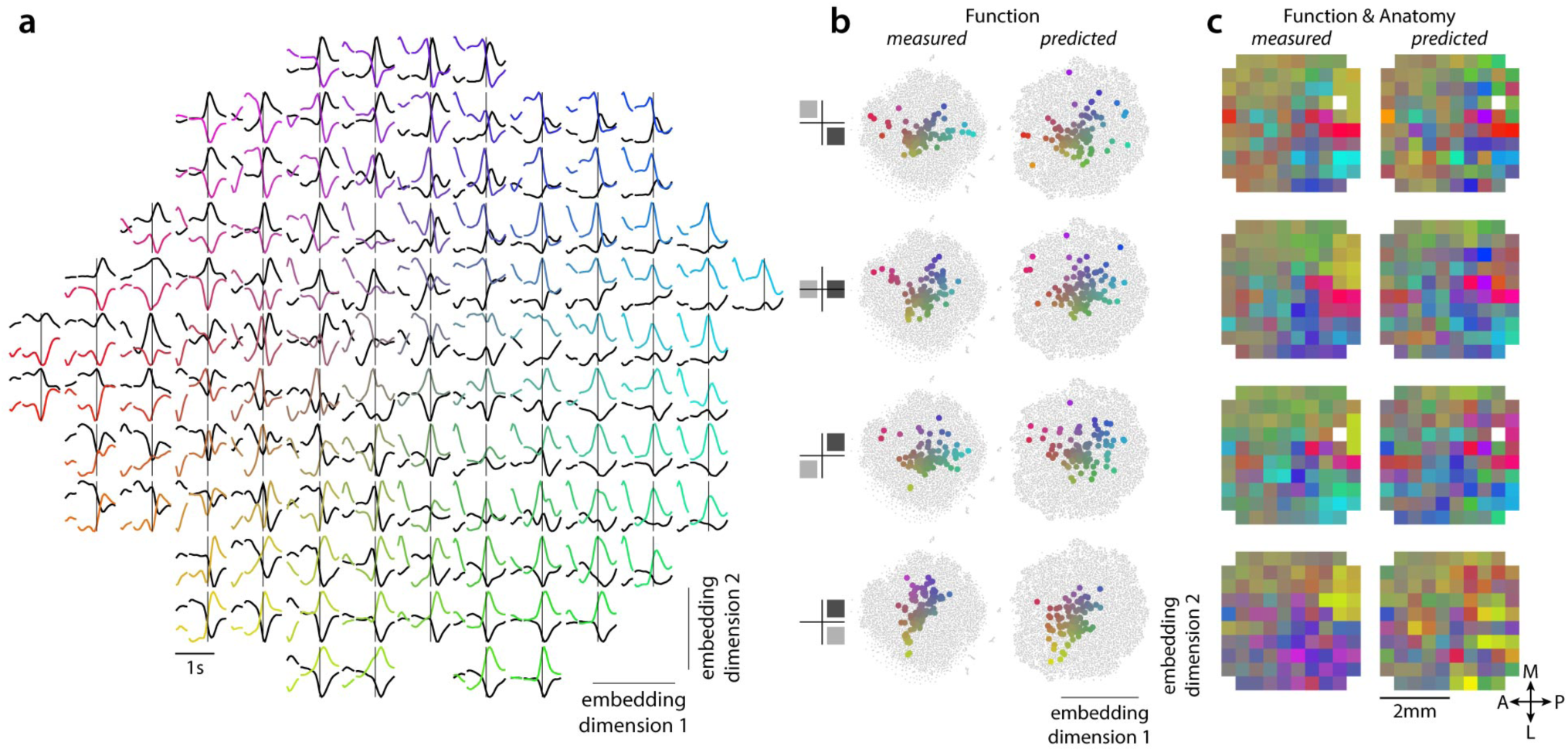
Response-field based predictions of dots-task responses in monkey V. Same conventions as in Fig. 7. **a**, Average unit responses based for the predicted responses, obtained as in Supp. Fig. 7b. **b**, Effect of cortical location and task-configuration on the unit responses observed in the dots-task. **c**, Measured and predicted topography of unit responses across the cortical surface. Each square corresponds to a circle in **b**, replotted with the same color at the corresponding array location.

**Supplementary Figure 9.**
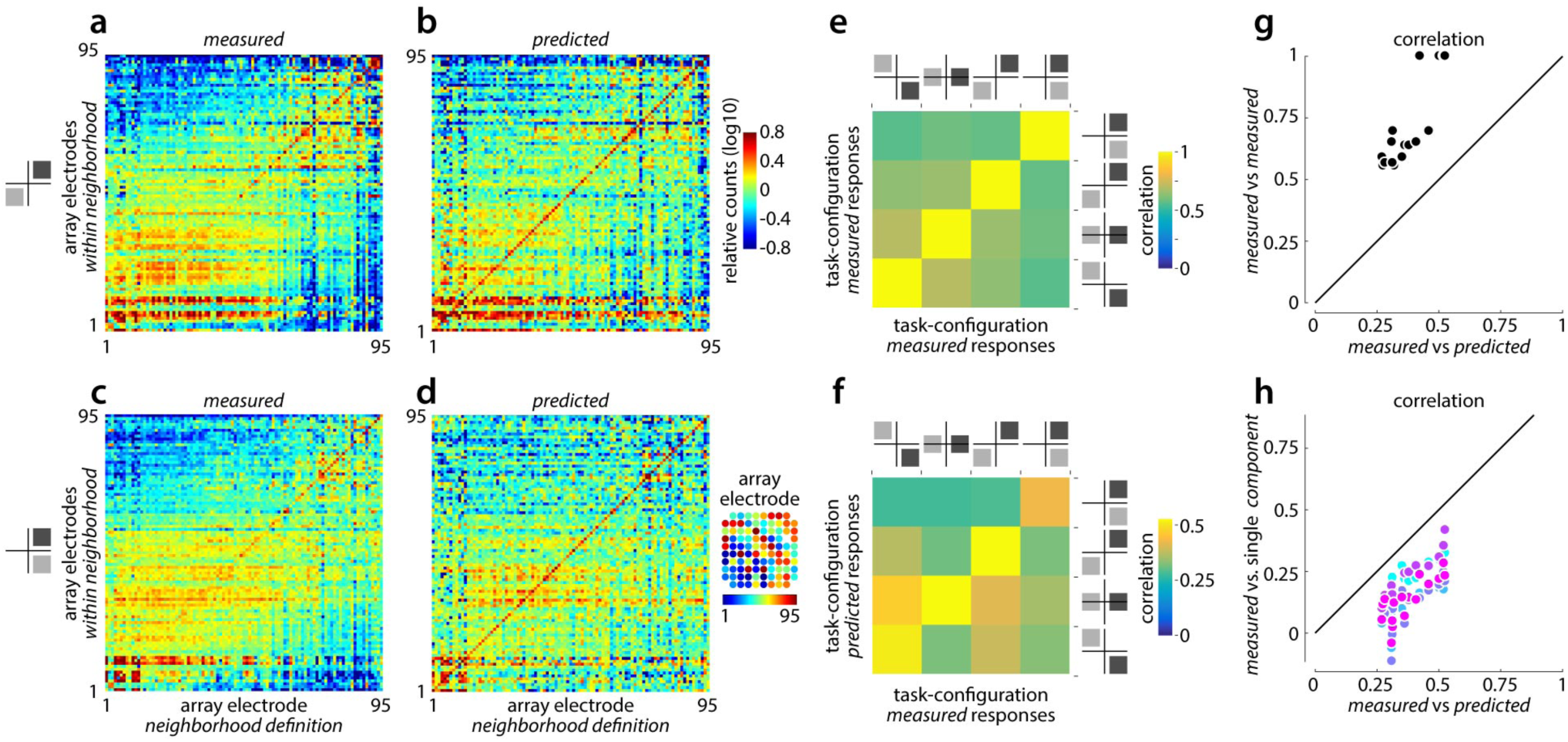
Accuracy of response-field based predictions in monkey V. Same conventions as in Fig. 8. **a**, Mixing-matrix for measured dots-task responses (monkey V) in one task-configuration (inset, left), with respect to array electrodes. **b**, Same as **a**, for responses predicted based on the response-fields estimated in the delayed-saccade task (Supp. Fig. 8). Same task-configuration as in **a**. **c**, Same as **a**, for a different task-configuration. **d**, Same as **b**, for the task-configuration in **c**. **e**, Correlation between measured mixing matrices from different task-configurations. **f**, Correlation between measured (horizontal axis) and predicted (vertical axis) mixing matrices from different task configurations, analogous to **e**. The predictions are most accurate when they are based on the same task-configuration as the measured responses (positive diagonal). **g**, Direct comparison of the correlations in **e** (vertical axis) and **f** (horizontal axis). **h**, The correlations in **f** (horizontal axis), for predictions based on all five choice-components, compared to correlations for predictions based on a single choice component (vertical axis). No single choice-component can account for the accuracy of the predictions for all task-configurations.

